# What Do Generative Models Learn About Adaptive Immune Receptor Repertoires? A Benchmark Study

**DOI:** 10.64898/2026.07.10.737788

**Authors:** Charlotte Würtzen, Maria Mamica, Chakravarthi Kanduri, Milena Pavlović, Victor Greiff, Bjoern Peters, Geir Kjetil Sandve

## Abstract

Generative models are increasingly used to model adaptive immune receptor repertoire (AIRR) sequence distributions, promising to decode the sequence diversity shaping immune responses and accelerate the design of therapeutic antibodies and T-cell receptors. Yet it remains unclear whether these models produce biologically meaningful outputs or merely capture surface-level sequence statistics while missing features driven by receptor generation and selection. Rigorous evaluation is needed, but the field lacks established standards, as existing machine learning metrics do not all translate directly to the AIRR domain, given the complex structure of the data and the lack of biological ground truth. Consequently, researchers face difficulties in evaluating the models and selecting appropriate ones, which can critically affect downstream clinical applications.

Here, we apply a suite of evaluation metrics tailored to AIRR sequence data and present a systematic comparison of popular generative model families proposed for the AIRR field, including variational autoencoders, long short-term memory networks, antibody language models, selection models, and simple statistical baselines. We focus specifically on the task of learning individual-specific immune receptor repertoires, a clinically relevant challenge with direct implications for personalized immunotherapy, disease monitoring, and vaccine response studies. By analyzing the sequences generated by each model, we identify memorization risks, innovation capabilities, and sensitivity to hyperparameter tuning. Taken together, these results advance the understanding of how current generative models reproduce the biology of individual immune repertoires and lay the groundwork for more principled model development and evaluation.

## Introduction

B-cell receptors (BCR, including antibodies) and T-cell receptors (TCRs), collectively called adaptive immune receptors (AIR), are components of the adaptive immune system that enable the recognition of foreign antigens with high specificity (1). This high specificity stems from an immense sequence diversity of TCRs and BCRs that is generated through V(D)J recombination, a somatic DNA rearrangement process occurring in the development of the lymphocytes, and by somatic hypermutation in the affinity maturation process for BCRs (2). Studying the characteristics of AIRR holds great potential to improve the diagnosis, treatment, and prevention of immune-related disorders (1,3). This has motivated the development of numerous computational approaches to understand the characteristics of AIR repertoires, including the use of generative models (1,4).

The application of generative models to study AIR sequences addresses several objectives, including density estimation of sequence probabilities (5–8), representation learning for downstream analyses (9–12), and, of particular interest in this study, the ability to sample realistic receptor sequences (13–16), with multiple methods supporting several of these objectives simultaneously (7,9,11). Methods capable of sampling receptor sequences employ a variety of generative architectures, including probabilistic recombination models (5–8,17), variational autoencoders (VAEs) (10,12,13), autoregressive models (11,14,15), generative adversarial networks (18,19), and, more recently, diffusion (20) and large language models (16,21). Correspondingly, sampling objectives vary from study to study and span simulating AIR(R) sequence libraries (16,18,21), generating individual (7) and cohort-level repertoires (8,13), and producing candidate antigen-binding receptors (14,15,20). As a result of this wide range of sampling motivations, the modeling scenarios considered in these studies differ notably, and are often described at varying levels of detail. Consequently, the choice of train and test datasets, along with preprocessing, and evaluation protocols vary widely, hindering the direct comparison between methods. This renders it difficult to select models for specific applications. To address this challenge, in this study, we evaluate whether generative models can reliably learn individual-specific immune repertoires’ distributions.

In the field of generative modeling, there exists no “gold standard” for method evaluation. However, Murphy outlined three general aspects to guide the assessment of generated samples (here defined as collections of generated instances): quality, diversity, and generalization (22). These aspects are defined relative to the target (original) distribution, which in this study refers to the true underlying repertoire distribution from which both the training and test subrepertoires are drawn (Figure 1c). Quality measures whether generated instances belong to the target distribution, diversity reflects the coverage of the target distribution, and generalization assesses the ability to go beyond the training data (22). Complementary to generalization is the concept of memorization, which refers to a model’s tendency to reproduce training samples (23,24). Although a variety of metrics have been proposed for generative model evaluation (22,25–31), the current generic metrics fail to adequately reflect the wide range of biologically relevant features present in diverse, high-dimensional AIR datasets, such as complex positional dependencies or network structures of repertoires (32–37). Furthermore, in some studies, assessments include metrics that rely on downstream predictions based on external representations or structure models (21,38–40), implicitly assuming that these models provide unbiased and reliable predictions. However, these assumptions are nontrivial for AIR sequence data, as available reference models are typically validated on conventional proteins, and it remains unclear whether their performance transfers to AIR sequences.

**Figure 1:**
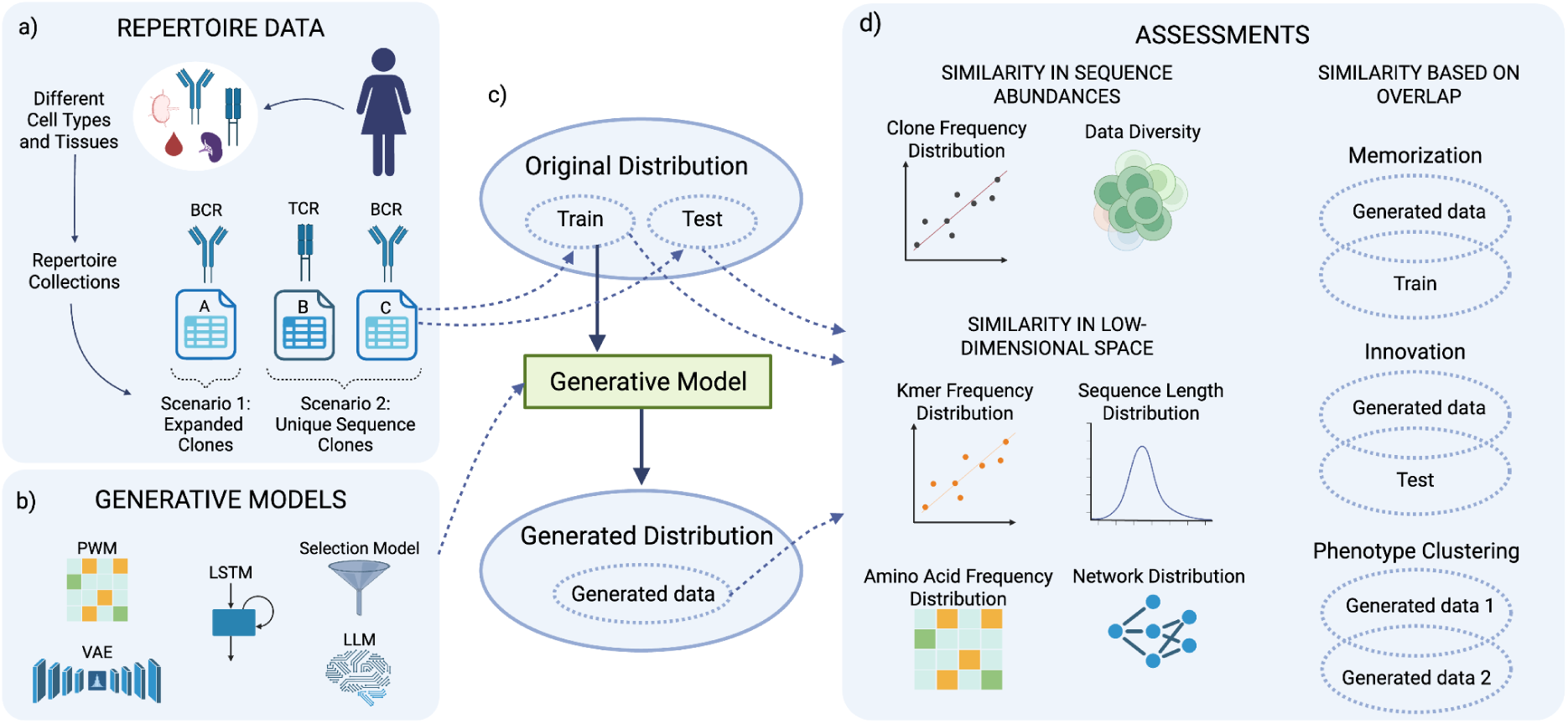
Overview of the benchmarking workflow assessing what generative models learn about AIRRs. a) Collections of TCR or BCR immune repertoires of different cell subsets and tissues were used to test models in two separate scenarios: 1) clonally expanded repertoires (collection A), and 2) unique clones repertoires (collection B and C). b) Studied generative model families: VAE, LSTM, the selection model SoNNia, the large language model ProGen2-OAS, and the baseline model PWM. c) Train and test sequences were sampled from collection repertoires (original distribution), whereof train samples were used to fit the generative models and the test samples held out for assessments. Subsequently, sequences were generated from the trained models. d) Dataset-based assessments of generative models comparing train, test, and generated data along three axes: similarity in sequence abundances (clone frequency distribution and data diversity), similarity in low dimensionality (k-mer, sequence length, amino acid, and network distributions), and similarity based on overlap (memorization against the train set, innovation against the test set, and phenotype clustering between generated sets), with each assessment applied to the relevant collections.

In this study, we selected and tailored a suite of evaluation metrics to explicitly capture relevant characteristics of AIRR sequences, with each metric addressing a distinct aspect of interest. We then applied them to profile diverse families of generative AIR(R) models, focusing on the realistic AIRR sequence generation objective. We profile variational autoencoders, recurrent neural networks (represented here by long short-term memory networks (LSTMs)), a targeted protein language model (ProGen2-OAS), along with domain-specific selection models (soNNia) (8,13–16). Together, these represent methodological diversity, spanning general-purpose to domain-specific models, while also reflecting some of the most commonly used methods in the AIR(R) field. We restrict our profiling to sequence-based models due to the limited availability of experimentally resolved 3D structures, which constrains the training and evaluation of structure-informed methods. Through complementary evaluation metrics, we gain insights into the strengths and weaknesses of the different generative model architectures, which may guide model selection and the further development of generative models in the field. Our analyses inspect how well the models learn subject-specific repertoire characteristics, measured by distributional comparisons of sequence length, amino acid, and subsequence composition; similarity of the repertoires’ network structure; their ability to generalize beyond the known training samples; and whether they capture tissue- or cell-subset-specific signatures.

## Results

Various generative AIRR models have been proposed, but their modeling objectives, experimental setups, and evaluation strategies differ considerably. Moreover, problem definitions are often underspecified, with key assumptions and modeling aims remaining implicit. Consequently, this lack of clarity may obscure what is actually being optimized or evaluated, hindering direct comparisons and cumulative progress across the field.

### Modeling individual repertoires is the natural starting point for profiling generative AIRR models

In this study, we explicitly define the shared modeling objective as learning the sequence distribution of a single individual-specific repertoire, where generalization refers to accurately capturing the underlying sequence distribution of that individual’s repertoire. We propose this as a starting point for model assessment, revealing whether models can learn intra-individual sequence diversity as a foundational step towards understanding the inter-individual diversity. Single-repertoire modeling provides a well-defined canonical problem formulation. It isolates the challenge of modeling the data-generating process in a single individual, namely V(D)J recombination and selection, without conflating it with questions of what should be shared or learned across individuals. Extending to inter-individual generalization, by contrast, allows various formulations depending on the number of donors, their relatedness, and the biological signal of interest. We explored the chosen objective in two scenarios: 1) learning the natural structure of repertoires, meaning the sequence distribution with clonal counts based on the natural clonal expansion, and 2) learning the distribution of repertoires with only unique sequence clones, thereby removing the dominance of high-frequency clones with the aim of better capturing the full sequence diversity. In the first scenario, large sequence overlaps were inevitable between train and test samples due to the clonal expansion and power law distribution of immune receptors (41), hindering assessment of generalization to unseen sequences which was instead more thoroughly explored in the second scenario. In addition, the unique sequence setting is applicable for studying shared recombination outcomes, selection pressures specific to cell subsets, and repertoire diversity both within and across individuals, without the confounding effect of clonal expansion.

We evaluated model capabilities in both scenarios using the proposed profiling workflow shown in Figure 1. The workflow includes a series of analyses designed to address relevant characteristics of AIRR sequences. For each model, we tuned hyperparameters to optimize recovery of k-mer (subsequence of length k) frequency distribution and, in the unique clones scenario, we also tuned an additional version for high innovation score. Optimization of the two criteria represent distinct aspects of learning the underlying distribution: one focusing on preserving the low-dimensional characteristics of the full distribution and the other one focusing on generalization to its unseen parts. The tuning criteria form the basis of our model naming convention (further details in Methods *Generative models: Hyperparameters tuning* and Table S1). By profiling these models on the shared modeling objective, we provide a first comparison of different generative model families, taking a step toward clarifying the state of generative modeling in AIRR.

### The majority of tested generative models are not reproducing clonal expansion structure

In the first scenario, we aimed to explore how well the models learn and reproduce the sequence frequency distribution of natural repertoires. To do so, we used data from collection A, which consisted of six pairs of test and train BCR subrepertoires with sequence frequencies reflecting the clonal expansion (details in Methods *Dataset collections*). Specifically, we asked whether the models could reproduce the clone frequency distributions observed in the held-out test sample. To address this, for each subrepertoire, we compared clone frequencies between test and generated samples, explored repertoire diversities, and inspected the direct memorization of train sequences (Figure 2, Figure S1, Figure S2a) (details in Methods *Evaluation*).

**Figure 2:**
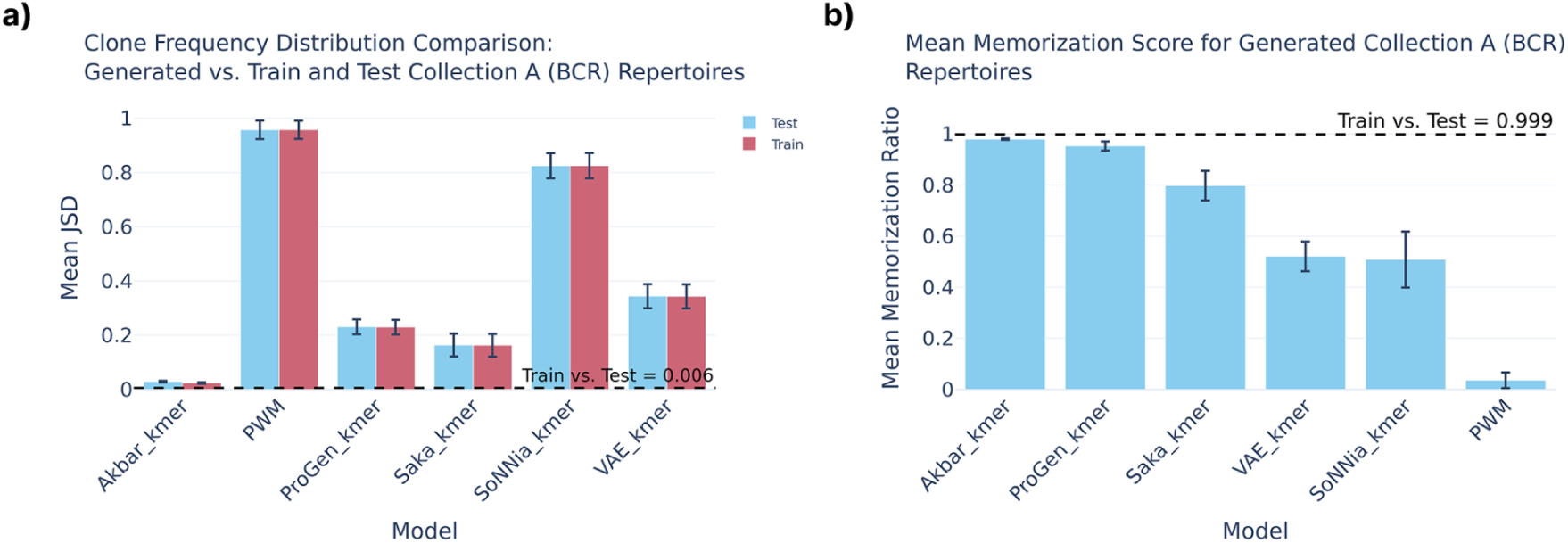
Assessment of models’ ability to learn the clone frequency distribution of clonally expanded repertoires (data collection A) revealed that most models are not able to reproduce the clonal expansion structure of the original repertoire. a) Mean Jensen-Shannon divergence scores between clone frequency distributions of generated and reference samples for each model. The horizontal dashed line indicates the mean reference JSD score between train and test clone frequency distributions. b) Memorization score per model averaged over generated subrepertoires. The dashed line indicates the reference score measured by the average overlap between paired train and test repertoires.

Large LSTM models (Akbars) reproduced clone frequency distributions most similar to those observed in the test repertoires, achieving a Jensen-Shannon Divergence (JSD) of 0.029, which was closest to the reference score between train and test repertoires (0.006) and substantially lower than all other models, which ranged from 0.163 (LSTM Saka) to 0.958 (Position Weight Matrix (PWM)) (Figure 2a). Visual inspection of clone frequency scatterplots confirmed these findings, with Akbar and Saka showing the densest concentration of points along the diagonal, while PWM and soNNia performed worst with highly scattered clones (Figure S1). The three worst-performing models (PWM, soNNia, and VAE) also exhibited the largest Shannon entropies, meaning they generated more diverse repertoires rather than reproducing the clonal expansion structure (Figure 2, Figure S2). Contrary to this pattern, Akbar’s diversities closely matched that of the test and train subrepertoires (Figure S2a). Furthermore, Akbar achieving best scores aligned with it generating the largest proportion of sequences directly memorized from the train samples (up to 93%) (Figure 2). Together, these results suggest that most of the evaluated models were not designed to capture clonal expansion, and that reproducing the frequency structure of expanded repertoires effectively requires memorizing the high-frequency clones seen during training. The architectural reasons underlying each model’s behavior are discussed in Supplementary Notes (“Model-specific factors limiting clone frequency recovery” section).

### Parameter-heavy LSTMs are prone to memorization even when trained on unique sequences

In a second scenario, we focused only on unique clones, which allowed us to better assess the models’ ability to generalize to unseen parts of the underlying repertoires. We fitted the models on 11 individual-specific TCR subrepertoires from collection B and 7 BCR subrepertoires from collection C, each paired with a matching same-size and non-overlapping test set from the same individual (details in Methods *Dataset collections*). While the high memorization rate observed for parameter-heavy LSTM models (Akbars) facilitated learning valid clone frequencies in the previous scenario, this had adverse effects when modeling the diversity of repertoires with unique sequences. The Akbar overlap- and k-mer-tuned models memorized on average 18.77% and 87.25% of train sequences for TCR respectively, compared to 55.21% and 84.93% for BCR (Figure S3). The remaining models exhibited rather low average levels of memorization, ranging from 0.06% to 5.18% for TCR repertoires and 0.01% to 9.2% for BCR repertoires. Although some degree of generated overlap with train sequences could be a natural consequence of learning the underlying distribution, these sequences were not informative for model assessments beyond memorization quantification and therefore subsequently replaced in a resampling step.

### Large language model achieves higher innovation scores, but the innovation is limited to small modifications relative to training data

To evaluate generalization, we measured the models’ ability to generate biologically plausible sequences not seen during training. Here and in further analyses, we were evaluating models using generated repertoires containing only novel sequences not overlapping with train subrepertoires (see Methods *Generative models: Training and sampling*). We quantified generalization through the innovation metric measuring the proportion of unique test sequences covered by the generative model. The innovation measure was restricted to unique generated sequences, preventing repeated sequences from inflating the scores (details in Methods *Evaluation: Innovation*).

When trained on unique TCR sequences, ProGen_overlap obtained the highest innovation scores (Figure 3a), with 373 to 700 unique innovative sequences overlapping with the test subrepertoires (1.73-3.26% of test sequences covered). Although ProGen generally produced lower clonal diversities and fewer unique sequences, it still covered the most sequences in the test set (Figure 3a, Figure S2). However, high innovation does not necessarily imply that ProGen covered the test repertoires with diverse innovative sequences or captured sequences substantially dissimilar from train sequences. Capturing diverse innovative sequences further from train instances demonstrates stronger generalization by learning the broader underlying structure of the distribution rather than simply staying in the vicinity of what is already known. To quantify the strength of the models’ innovative power, we measured the Hamming distance of innovative sequences to their closest train sequences. In general, innovative sequences showed high similarity to train sequences for all generative models fitted on TCR repertoires, with most sequences only differing by Hamming distance 1 or 2, and no sequences differing by more than distance 3 (Figure 3c). This shows that for all models, including the large language model ProGen, innovative capabilities were only found within a close range to the known train sequences. This pattern was further pronounced for both ProGen models when considering the full generated repertoires rather than innovative sequences only (Figure 3d). Compared to the reference distance distribution of test sequences, ProGen generated fewer sequences with greater distance to train, suggesting that ProGen generally explored a more limited region of the sequence space.

**Figure 3:**
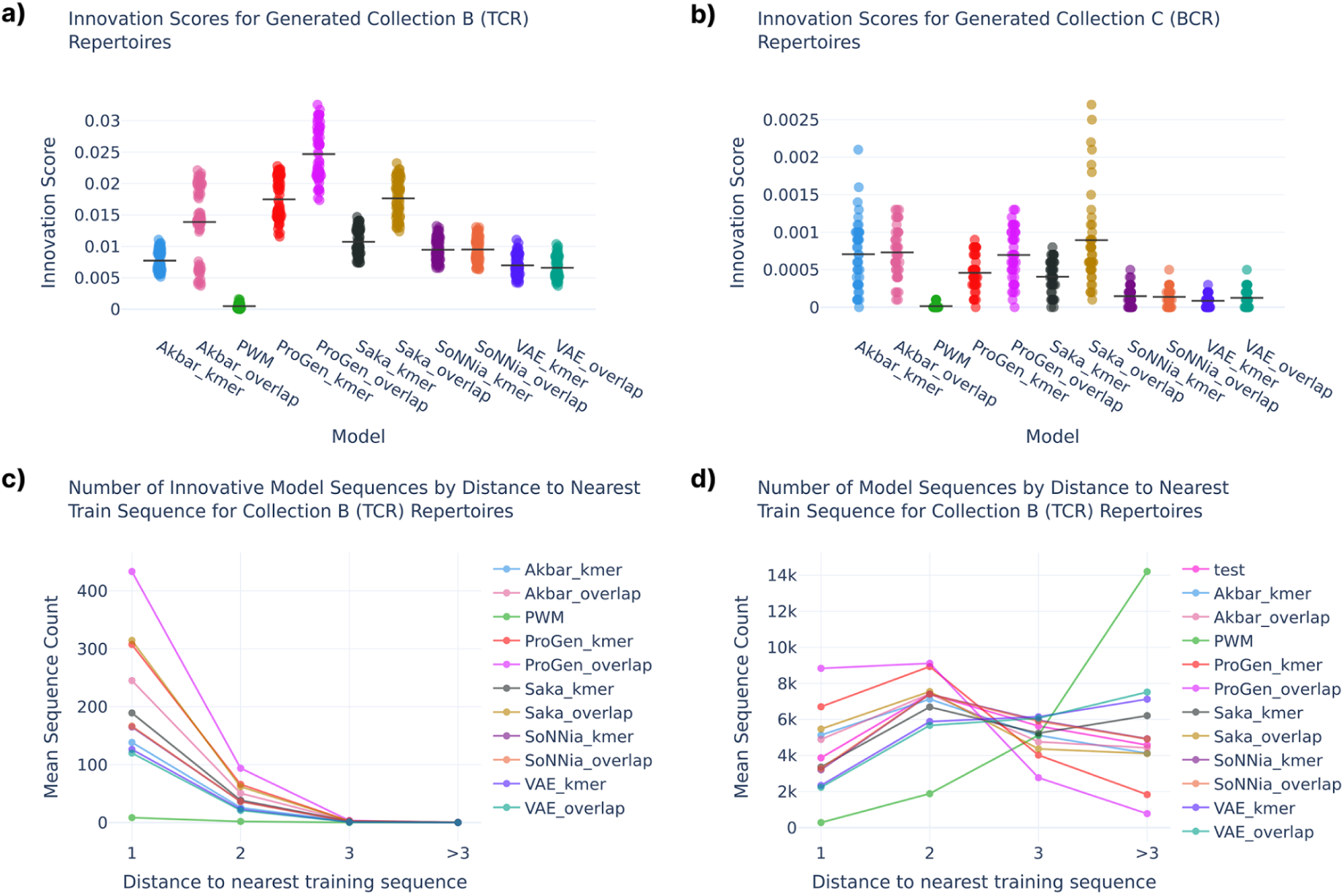
Analysis of innovative capabilities of generative models revealed that ProGen produced the highest local innovation. a,b) Innovation scores per generative model for TCR and BCR repertoires, respectively. Each dot represents a generated repertoire, and lines mark the average innovation scores over repertoires per model. c) Number of innovative sequences within different Hamming distances from train sequences. Sequence counts are averaged over repertoires and dataset splits. d) Average number of test or generated sequences (full samples) within different Hamming distances from train.

For BCR repertoires, the number of innovative sequences was generally lower than for TCR (ranging 0–27). This may be explained by the smaller dataset sizes (10,000 vs. 21,500 sequences) as well as the tendency for CDR3 sequences to be longer for BCRs (Figure S4), both of which reduce the likelihood of finding direct overlap with test repertoires. For the generated BCR repertoires, it was the Saka_overlap model which showed highest innovation scores (Figure 3b). However, these high scores of Saka_overlap were all found for 6 data splits originating from the same model trained on one repertoire (6279_pLN_BCR), which also exhibited inconsistent behavior compared to other repertoires in terms of clonal diversities (Table S2, Figure S2d). Given this, together with the generally low range of innovative sequences for BCR, it is difficult to draw any claims about Saka’s advantage over other models. Notably, ProGen did not show a clear advantage on BCR repertoires despite being pre-trained on BCR sequences. These results could be evidence that fine-tuning the large ProGen requires having more train sequences than in the BCR datasets used in our study.

### Models learn connectivity distributions better when tuned to preserve low-dimensional characteristics than when tuned for sequence innovation

We evaluated overall distributional similarities between generated and original repertoires across multiple dimensions, including reduction of the high-dimensional amino acid CDR3 sequences to length, positional amino acid frequency, and k-mer frequency distributions. Additionally, we compared connectivity distributions of repertoires which were computed from the sequences’ Levenshtein-based neighbor counts (see Methods *Evaluation: Data consistency in low-dimensions: amino acid, sequence length, k-mer, and connectivity distributions*).

Figure 4 shows the distributional comparisons across the tested models, quantified by JSD scores, which revealed that generative model performance was highly sensitive to the choices made in hyperparameter tuning. Models tuned for k-mer recovery tended to generally achieve better distributional similarity compared to the same models tuned for high innovation score, with only two exceptions (Saka for collection B and VAE for collection C) showing opposite behavior in length distributions (Figure S4a,b). This was expected for k-mer frequency distributions, but the same advantage extended to connectivity distributions (Figure 4e,f) and to positional amino acid frequencies (for most cases, see Figure S5), neither of which was directly optimized during tuning. Generally, the large language and LSTM-based models showed stronger sensitivity to tuning criteria compared to the VAE and soNNia models.

**Figure 4:**
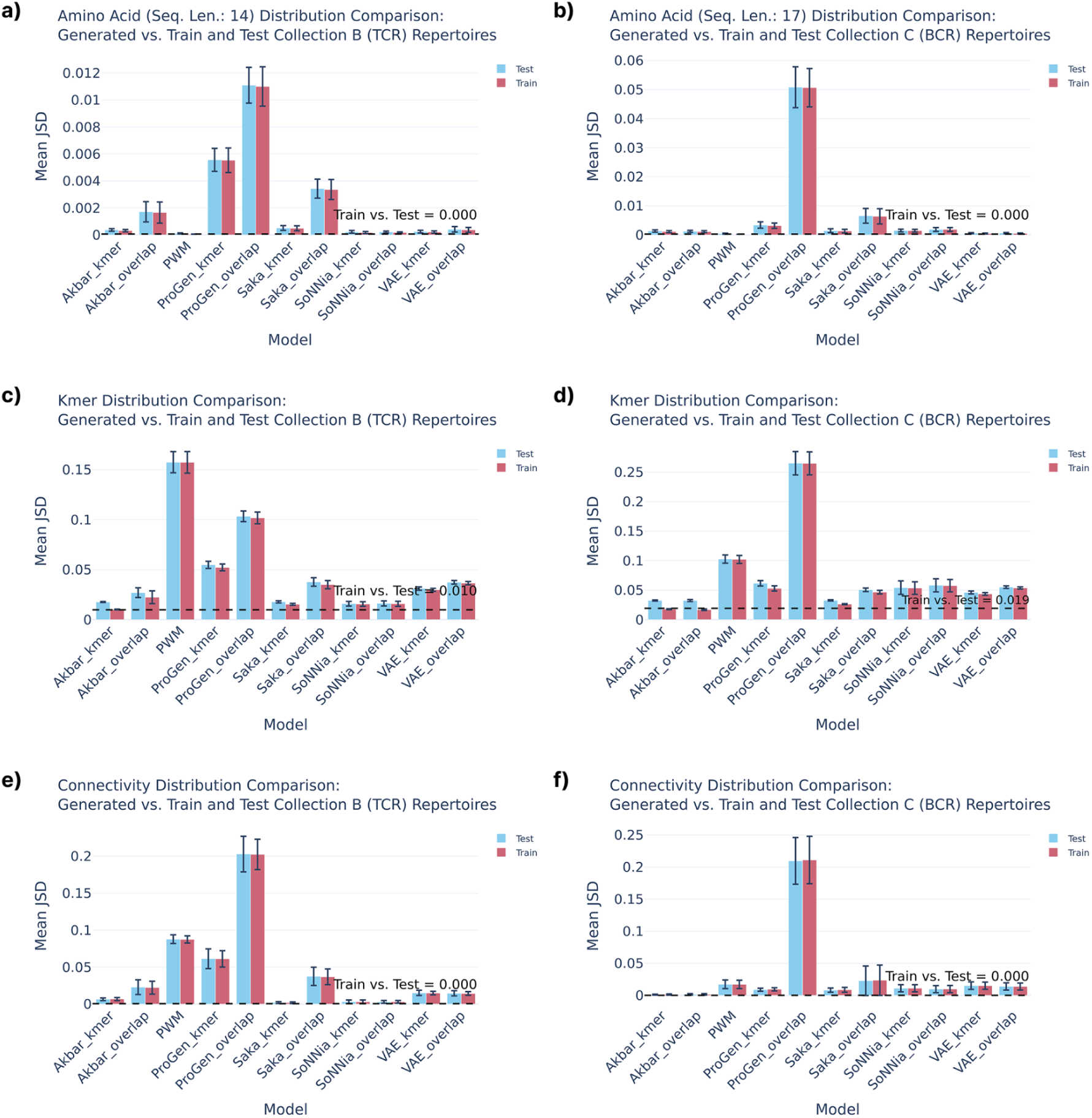
Models tuned to preserve k-mer recovery tended to achieve lower Jensen-Shannon divergence scores across all lower-dimensional metrics than those tuned for innovation. Models were evaluated by JSD scores for train, test, and generated TCR and BCR sequences (unique clones scenario), comparing sequence-level properties across models and tuning strategies. The sequence distributions were compared across (a,b) positional amino acid frequencies of middle length sequences (14 for TCR and 17 for BCR), (c,d) k-mer frequencies, and (e,f) neighbor counts (connectivity) of unique sequences with neighbor defined as another sequence with Levenshtein distance of 1. A lower JSD indicates higher distributional similarity. For each model, the average JSD between train and generated, and test and generated was computed from the subsets of generated data. All dashed lines show corresponding reference scores computed between train and test sets.

Overall, ProGen (tuned for innovation) exhibited the worst recovery of low-dimensional characteristics. We therefore examined repertoire-specific connectivity distributions, which showed that ProGen_overlap tended to produce sequences with more neighbors within a Levensthein distance of 1 (Figure S6c). This higher repertoire connectivity compared to reference subrepertoires is evidence that ProGen_overlap learned to generate more dense sequence regions, causing it to also obtain poorer divergence scores across the other reduced dimensions. For both the large language and LSTM models, the optimized temperature hyperparameter was lower for innovation tuned vs. k-mer tuned models (0.8 vs. 1.0, see Table S3), narrowing the generated distribution and reducing the sampling diversity. Potentially, models tuned for high innovation could be focusing on more dense local areas rather than covering the full space of a natural repertoire’s sequence distribution. In these dense regions, such models are more likely to discover a sequence matching the test set, boosting the innovation score by oversampling sequences close to the train data.

The shown effects of tuning indicate that optimising the model for one purpose, such as discovering novel valid sequences, may result in worse performance on other biologically relevant metrics. In profiling of generative model families, it becomes ambiguous which version of a method should be used for comparison and underlines the importance of clearly defined experimental setups and objectives.

### Large language models are able to learn cell-subset-, and subject-specific characteristics

As an additional task, we examined whether the models were able to capture repertoire-specific patterns related to tissue, cell-subset, or to the given subject. For each generative method, we fitted one model per repertoire, separately for the repertoires from collection B (labeled by subject and cell type) and collection C (labeled by subject and tissue). Within each method-collection combination, every fitted model was then used to generate a synthetic repertoire, and the resulting repertoires were clustered based on pairwise Jaccard similarity. Finally, we assessed whether the recovered clusters corresponded to the labels of the original training repertoires (Figure 5a). In addition, we computed Mean Average Precision (MAP) as rank scores, separately for label and subject (see Figure 5a and Methods *Evaluation: Learning cell-subset-, tissue-, and subject-specific patterns*).

**Figure 5:**
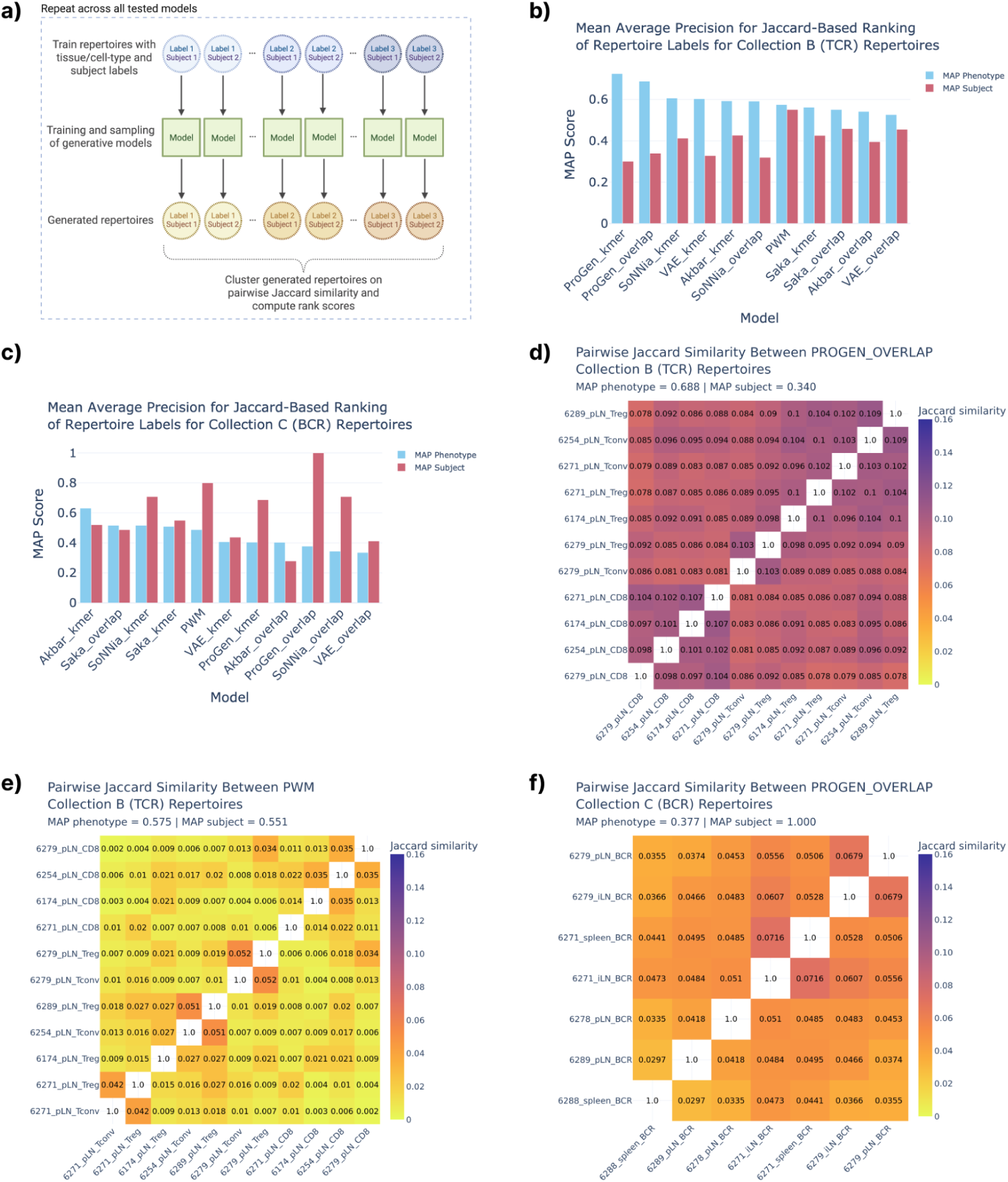
Hierarchical clustering of pairwise Jaccard similarity scores between generated repertoires demonstrated that ProGen was able to learn both cell-subset- and subject-specific characteristics. a) Overview of the experimental setup. Train repertoires from identical and distinct tissue, cell-subset, and subject were used to fit separate instances of the same generative model, from which synthetic sequences were sampled. Generated repertoires were then clustered based on pairwise Jaccard similarity to assess whether repertoires generated from the same label or subject clustered together. b) MAP scores per model for TCR repertoires. c) MAP scores per model for BCR repertoires. d) ProGen_overlap fitted on TCR repertoires from CD4+ T conventional, CD4+ T regulatory, or CD8+ T-cells subsets. e) PWM fitted on the same TCR repertoires as in d. f) ProGen_overlap fitted on BCR repertoires from spleen, irrelevant lymph node (ILN), and pancreatic lymph node (PLN) tissues.

For TCR repertoires, both ProGen models achieved the highest MAP scores for the cell-subset ranking, suggesting that they best captured cell-subset-specific sequence patterns among all evaluated models (Figure 5b). This was further supported by the clustering analysis, where CD8+ repertoires clustered together (Figure 5d, Figure S7). Clustering was less apparent for T conventional and T regulatory cell subsets, though in repertoires generated by ProGen optimized for innovation, some subject-specific structure was visible (subject 6279 and to a lesser extent subject 6271). However, this was not reflected in the MAP scores for subjects for ProGen, suggesting that the model failed to consistently capture subject-specific patterns and predominantly captured cell-subset patterns instead. Interestingly, when considering MAP scores for subject, the simplest model PWM achieved the highest scores, which was also supported by the heatmap in Figure 5e.

For BCR repertoires, no model was able to cluster the same tissues together (Figure 5f, Figure S8), which may suggest that tissue-specific differences are too subtle to be captured by the models, or that the repertoire sizes were insufficient (60,000 for BCR vs. 129,000 for TCR). Although tissue-specific patterns were not recovered, ProGen_overlap achieved a perfect MAP score of 1 for individual ranking, meaning that it correctly separated subjects (Figure 5c, Figure 5f). This should however be interpreted carefully, as the BCR collection consisted of only 7 repertoires in total, with two subjects represented by two repertoires each and three subjects represented by a single repertoire.

## Discussion

Although a growing number of generative models have been proposed for AIR(R), they have been assessed under widely varying experimental conditions, hindering their direct comparison. To address this, we evaluated representative models of multiple generative architecture families on a shared modeling objective: learning the sequence distribution of an individual-specific AIRR. To the best of our knowledge, this constitutes the first systematic comparison of generative models for AIRR sequences.

So far, generalization in AIRR generative modeling has typically been tested indirectly through held-out sequence probability estimation (8,13,42) or proxies like sequence novelty or diversity relative to the training set (14–16). However, generalization has rarely been explicitly and consistently defined in the generative setting and the term is largely absent from the existing literature, underscoring the need for more standardized and comprehensive evaluation frameworks (33,43). Here, we explored generalization by the sequence overlap between generated and held-out test subrepertoires as a measure of whether the models could produce novel yet plausible sequences. Our analyses revealed a notable limit on generalization: although the antibody-specific language model captured most unseen test sequences, this was only achieved by exploiting the ability to learn close neighborhoods of train sequences. In fact, none of the generative methods discovered the test sequences with higher distance (≥3) to train sequences. Thus, while the models achieved a form of innovation, its local nature highlights that more precise conceptualization of innovation remains an open challenge. Key questions include whether innovation is a binary property or a matter of degree, and whether the observed innovation stems from learning the full underlying distribution or by concentrating on denser, easier-to-model regions of sequence space.

Memorization, in turn, may be a natural consequence of learning the distribution, and sometimes even helpful as seen in LSTM’s ability to reproduce clone frequencies of clonally expanded repertoires. In the context of innovation and generalization, however, high sequence memorization shifts from a feature to a limitation. While we introduced a definition as direct sequence reproduction, memorization may also occur in the vicinity of train sequences, leaving room to further study the relationship between memorization and generalization. In the broader context of generative modeling, notions such as memorization (23,24,44) and novelty (45,46) have already been explored as proxies for the overfitting and generalization concepts; however their definitions and interpretations differ across studies.

Another aspect explored in this study was whether the models are able to capture repertoire patterns arising from tissue, cell subset, or subject characteristics. We found that ProGen was able to capture cell-subset-specific patterns for TCRs as well as subject-specific patterns for BCRs. These results point toward the potential utility of generative models in discriminative tasks leveraging repertoire-level distributional differences. As discussed by Tomczak (47), generative models may be particularly useful in settings where decision-making benefits from uncertainty quantification. Furthermore, the findings suggest the potential in applying generative models to study biological structure arising from convergent recombination, subset-specific selection, or intra- and interindividual variability (48).

This study further underlined that highly parameterized models are particularly sensitive to which criterion is optimized during hyperparameter tuning. This is consistent with the broader observation that generative model evaluation criteria can be largely independent of one another, as a model that excels by one measure may fall short by another (49). Some hyperparameters might control a trade-off between competing criteria rather than optimizing one. Sampling temperature is the canonical example: a single trained model behaves differently as temperature varies, trading fidelity against diversity (50). Ideally, when profiling generative models, they should be compared across multiple tuning criteria and evaluation metrics, though in practice this is limited by the computational cost of training generative models (see Table S4 and Table S5 for a computational overview of all tuning and training runs in this study). Furthermore, when reproducing existing methods or developing new ones, researchers should carefully align their choice of tuning criterion with their intended research goals, as this can significantly shape model behavior. They should also be careful about making claims on (contrastive) characteristics of models based on exploration of only a single configuration, since any such conclusions may be highly sensitive to what was selected as objective for hyperparameter tuning.

In this work, we compiled a curated set of evaluation metrics suited to the AIRR domain, covering both statistical sequence properties and biologically relevant repertoire characteristics. These can guide method developers in designing their evaluation protocols, as well as inform further benchmarking efforts in the field. They may also serve as an inspiration for researchers working beyond the AIRR domain, including the broader area of protein engineering, given the shared nature of sequence data. Furthermore, as the modeling objectives for AIRR generation are diverse, we encourage future works to be explicit about chosen objectives and experimental setups, as this is a prerequisite for meaningful cross-study comparison. Finally, we highlight the need for further assessment studies in the AIRR field. As the number of generative methods continues to grow, systematic benchmarking across diverse modeling scenarios, such as antigen-specific binder generation or population-level repertoire modeling, will be essential for identifying methodological progress and guiding model selection for downstream biological and clinical applications. With this study, we lay the groundwork for clearer understanding of the current state of generative modeling in AIRR as well as for more systematic evaluation practices for a broader variety of modeling scenarios in AIRR and beyond.

## Methods

### Dataset collections

To compare the generative models, we prepared three different collections of TCR and BCR repertoires, all retrieved from the iReceptor database (51). In this work, a repertoire refers to the sample of CDR3 sequences from a single individual, and a subrepertoire refers to a subsample of that repertoire. We explored two distinct modeling scenarios: 1) repertoires with CDR3 sequences of clonally expanded cells and 2) repertoires with only unique CDR3 sequences retained. In both scenarios, sequences were sampled without replacement (i.e. divided) into train and test subrepertoires of equal size, resulting in no sequence overlap between the train and test datasets in scenario 2.

- Collection A (scenario 1): BCR repertoires originated from a study by Sokal et al. (52). We weighted CDR3 sequences based on unique molecular identifier (UMI) counts for each of the repertoires, and sampled train and test subrepertoires of 150,000 sequences each (determined by the repertoire sizes), with highly expanded clones represented in both.
- Collection B (scenario 2): TCR repertoires originated from a study by Seay et al. (53), and were chosen due to higher numbers of unique CDR3 sequences than in collection A, as well as the study’s separation into repertoires of different cell subsets and tissues. This collection comprised TCRbeta donor-specific train and test datasets containing 21,500 sequences each. Repertoires consisted of either CD4+ conventional, CD4+ regulatory, or CD8+ TCR beta sequences.
- Collection C (scenario 2): BCR repertoires were obtained from the same study as for collection B to further explore the setting with unique clones only, this time on BCR heavy chain repertoires. As sequence counts were lower in these repertoires, 10,000 sequences were sampled for each train and test set. In this collection, the tissue specificity was now varied with repertoires coming from either the pancreatic lymph node or the spleen.

To consider the consistency of the methods and account for the evaluation scores’ variability across repertoires, we conducted several runs per model, each on a different repertoire from the given collection. However, as the need for computational resources can quickly escalate when comparing multiple different models fitted separately on many repertoires, we limited our analyses to only 7, 12, and 8 repertoires for the three collections, respectively, with one repertoire being held out from each collection for hyperparameter tuning. Table S6 shows an overview of the data collection details while a brief summary is given in Table S7.

### Generative models

In this work, we chose representatives from four different model families. Additionally, as a simple baseline, we fitted a Position Weight Matrix (PWM) model based on position-wise amino acid frequencies. The implementations of other models were either based on published implementations or inspired by published studies: here we fitted VAEs based on Davidsen et al. (13), LSTMs based on Akbar et al. and Saka et al. (14,15), linear versions of the SoNNia method (as recommended by authors for small dataset sizes) (8), together with large antibody-specific language models based on the original pretrained ProGen2-OAS (16) (referred to as ProGen), which was subsequently fine-tuned on our collections.

#### Hyperparameters tuning

To accommodate the multifaceted nature of generative model assessment and selection, we tuned hyperparameters of these models on held-out repertoires according to two different criteria (k-mer recovery and direct sequence overlap) for the unique clones scenario, and one criterion (k-mer recovery) for the clonally expanded scenario. This resulted in the selection of up to two models per family (excluding the baseline PWM). Using grid searches, we selected for each scenario one model configuration that performed best at recovering subsequences of length 3 (k-mers with k=3) frequency distributions. For the unique clones scenario, we additionally selected a model configuration that achieved the highest sequence overlap between generated data and the test sample (described later as innovation score). This criterion was not used for tuning in the clonal expansion scenario due to the low number of unique test sequences absent from the train (<79 depending on the repertoire). To distinguish between the two differently chosen models, we introduced the suffixes ‘_kmer’ and ‘_overlap’ as the naming convention. Table S3 shows the hyperparameters of the chosen models, whereas Table S5 summarizes the scale of the hyperparameter tuning experiments.

#### Training and sampling

All generative models were trained, and sequences for further assessments were sampled, using the open-source immuneML platform (54). Table S4 summarizes the scale of the computational experiments, including the numbers of trained models and sampled repertoires. For the unique clones scenario, memorized sequences were replaced by additional sampling from each model until the same sample size as train and test was reached (within-sample repetitions were allowed). Replacing memorized sequences allowed us to test whether a model learned repertoire characteristics generalizing beyond its training data. We therefore applied this re-sampling only after quantifying the memorization level.

### Evaluation

From each of the fitted models, we generated 450,000, 130,000, and 60,000 sequences when trained on data collections A, B, and C, respectively. For most analyses, besides the clustering experiment, assessment scores were computed on generated data split into batches matching training and test sample sizes. This resulted in three batches for collection A given its larger sequence count and computational cost, and six batches for collections B and C. To reduce sampling variability, scores were then averaged across batches. All evaluations were performed using CDR3 sequences only. For scenario 2 (unique clones), assessment was conducted based on generated repertoires in which sequences overlapping with train subrepertoires had been replaced by additional sampling (see Methods *Generative models: Training and sampling*), except for assessment of memorization ratios.

#### Clone frequencies

For collection A (scenario 1), we assessed the ability to reproduce the statistical structure of natural repertoires with clonal expansion. Jensen-Shannon divergence (JSD) scores (55) were computed between generated and test clone frequency distributions to compare model performances. JSD is defined as:

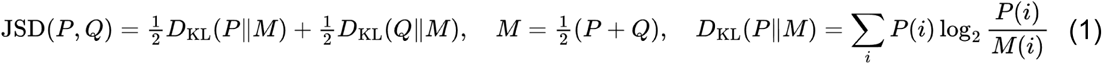

where *D_KL_* is the Kullback-Leibler divergence, and *P* and *Q* are the generated and test clone-frequency distributions over all clones observed in either sample. Using base-2 logarithms, JSD ranges from 0 (identical distributions) to 1 (disjoint), with lower values indicating closer agreement.

#### Clonal diversity

To analyse clonal diversity, for each repertoire we computed four measures based on CDR3 sequence counts: Shannon entropy (Equation 2), Pielou’s evenness (Equation 3), the Gini coefficient (Equation 4) and the Gini-Simpson Index (Equation 5) (56), defined as follows:

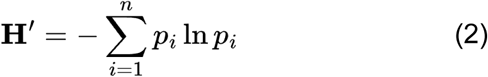

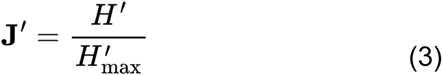

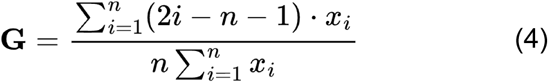

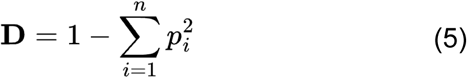

where p_i_ is the frequency of clone i, n is the total number of unique clones, H_max_ is the maximum possible Shannon entropy for a repertoire of n unique clones (Equation 3 only), and x_i_ are clone counts sorted in ascending order (Equation 4 only). For collections B and C, consisting of repertoires with only unique clones, the computed diversity reflected the theoretical maximum diversity scores and served as a reference for comparison with generated repertoires.

#### Memorization

We defined a memorization score as the proportion of generated sequences with a direct sequence overlap to the train sample used to fit the given generative model. The scores were computed using the *existence* command of the CompAIRR tool (57) before sampling additional sequences to obtain generated repertoires of only novel sequences.

#### Data consistency in low-dimensions: amino acid, sequence length, k-mer, and connectivity distributions

To assess the ability of generative models to reproduce central low-dimensional aspects of AIRR distributions, we computed the distributions of CDR3 sequence lengths, positional amino acid frequencies, k-mer frequencies, and sequence connectivity of each train, test, and generated repertoire. Using the JSD (Equation 1), we compared the distributions of sequences sampled from each fitted generative model to the corresponding train and test sequence distributions for each metric. The comparisons of positional amino acid frequency distributions were stratified by sequence lengths, exploring lengths ranging from 10-20, to avoid potential noise caused by sequence alignments and differences in sequence length distributions. In the analysis comparing k-mer frequencies, we investigated the frequency distributions of subsequences of length 3 (k=3). For connectivity, we measured the number of neighbors for each of the unique CDR3 sequences in the repertoire, where a neighbor was defined as a distinct sequence that differs by a Levenshtein distance of 1, at the amino acid level (58).

#### Innovation

We defined innovation as a model’s ability to generate unique sequences found in the test set but absent from the training set, indicating that the model has learned to sample from the underlying distribution beyond the observed training data. More concretely, we formulated the innovation score as the proportion of unique unseen test sequences that were covered by at least one direct sequence match in the generated data, drawing inspiration from recall formulations for the evaluation of generative models (27,28). We mainly explored innovation in the unique sequences scenario, while innovation for the clonal expansion scenario is discussed in Supplementary Notes (“Innovation in the clonal expansion scenario” section) as assessment was highly challenged by the small unique sequence overlap between train and test subrepertoires. To further study the innovation capabilities in the unique sequences scenario, we measured the Hamming distance of innovative sequences to their nearest train sequences, resulting in a distance distribution across values 1, 2, 3, and >3. Due to low numbers of innovative sequences in generated BCR repertoires (0-27), we limited the BCR evaluation to innovation scores only, omitting analysis of sequence distances to train. All sequence overlaps were computed using the *existence* command of the CompAIRR tool (57).

#### Learning cell-subset-, tissue-, and subject-specific patterns

To investigate whether the generative models captured patterns specific to tissue- or cell-subset repertoire labels or subject IDs, we measured similarities between generated repertoires from models trained on repertoires of different labels and individuals. This analysis was restricted to the unique clones scenario (collection B and C), as similarities between clonally expanded repertoires were dominated by memorization, which can reflect repertoire-specific patterns without genuine learning of such characteristics. In collection B, separate models were fitted to repertoires of different cell subsets: CD4+ T conventional, CD4+ T regulatory, and CD8+ T-cells. Based on collection C, we explored differences between tissue-specific cells from the spleen, pancreatic lymph node, and “irrelevant” nonpancreatic lymph node. For each generative method, we fitted one model per repertoire within a given collection and computed pairwise Jaccard similarities based on direct sequence overlap between unique generated sequence sets. Based on these similarities, we performed hierarchical clustering using average linkage. Furthermore, to quantify the degree to which generated repertoires were more similar to repertoires of the same label or subject, we computed Mean Average Precision (MAP) (59) scores based on the pairwise Jaccard similarity matrix. For each repertoire, all others were ranked by descending Jaccard similarity and Average Precision was computed over repertoires sharing the same label or subject, excluding self-comparisons. MAP was then calculated as the mean Average Precision across all repertoires, as follows:

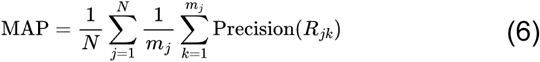

where N is the number of repertoires, m_j_ the number of repertoires sharing the same label or subject as repertoire j, and R_jk_ the set of repertoires ranked from the top down to the k-th relevant one.

## Acknowledgements

We thank Natalia Rincon for valuable discussions on a potential method to benchmark.

## Author Contributions

CW, MM, CK, BP, and GKS conceptualized the study. CW and MM performed data curation, designed and created software, and performed analyses. CW, MM, and GKS drafted the manuscript. CK, MP, VG, BP, and GKS provided critical feedback. CK, BP, and GKS supervised the project.

## Data and Code Availability

Experiments were done on publicly available data only. Repertoires of collection A were from the study by Sokal et al. (52) while repertoires of collection B and C were gathered from a study by Seay et al. (53). All data was retrieved from the iReceptor database (51). The code for all stages of the evaluation pipeline can be found at: https://github.com/sandvelab/gen-airr-profiling.

## Supplementary Material

## Supplementary Figures

**Figure S1:**
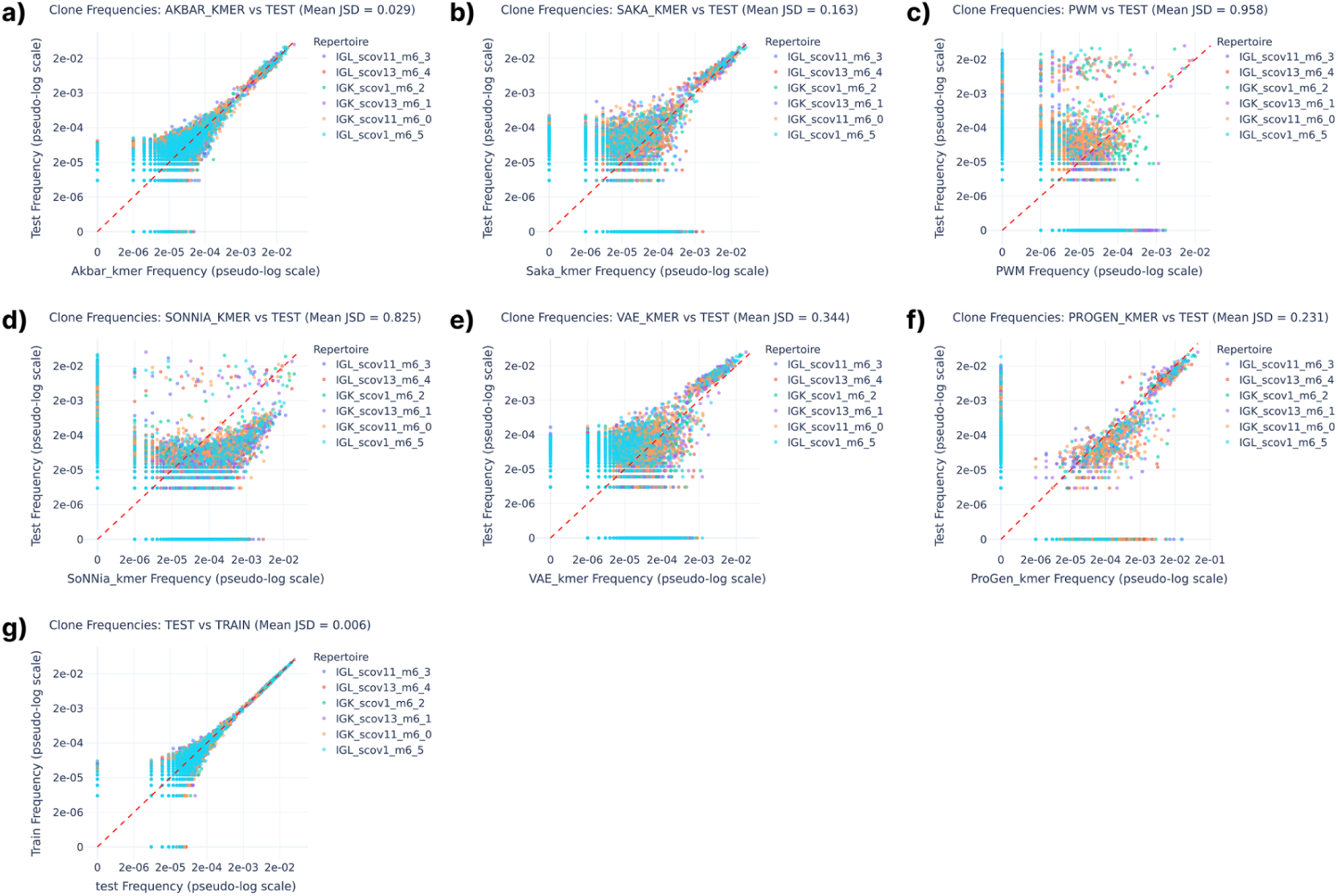
Assessment of models’ ability to learn the clone frequency distribution of clonally expanded repertoires (data collection A) revealed that most models are not able to reproduce the clonal expansion structure of the original repertoire. a-f) Scatterplots of clone frequencies for generated vs. test repertoire samples for models Akbar, Saka, PWM, soNNia, VAE, and ProGen, respectively. Diagonal reference lines indicate equal frequencies in the two samples. g) Scatterplot of clone frequencies for train vs. test samples.

**Figure S2:**
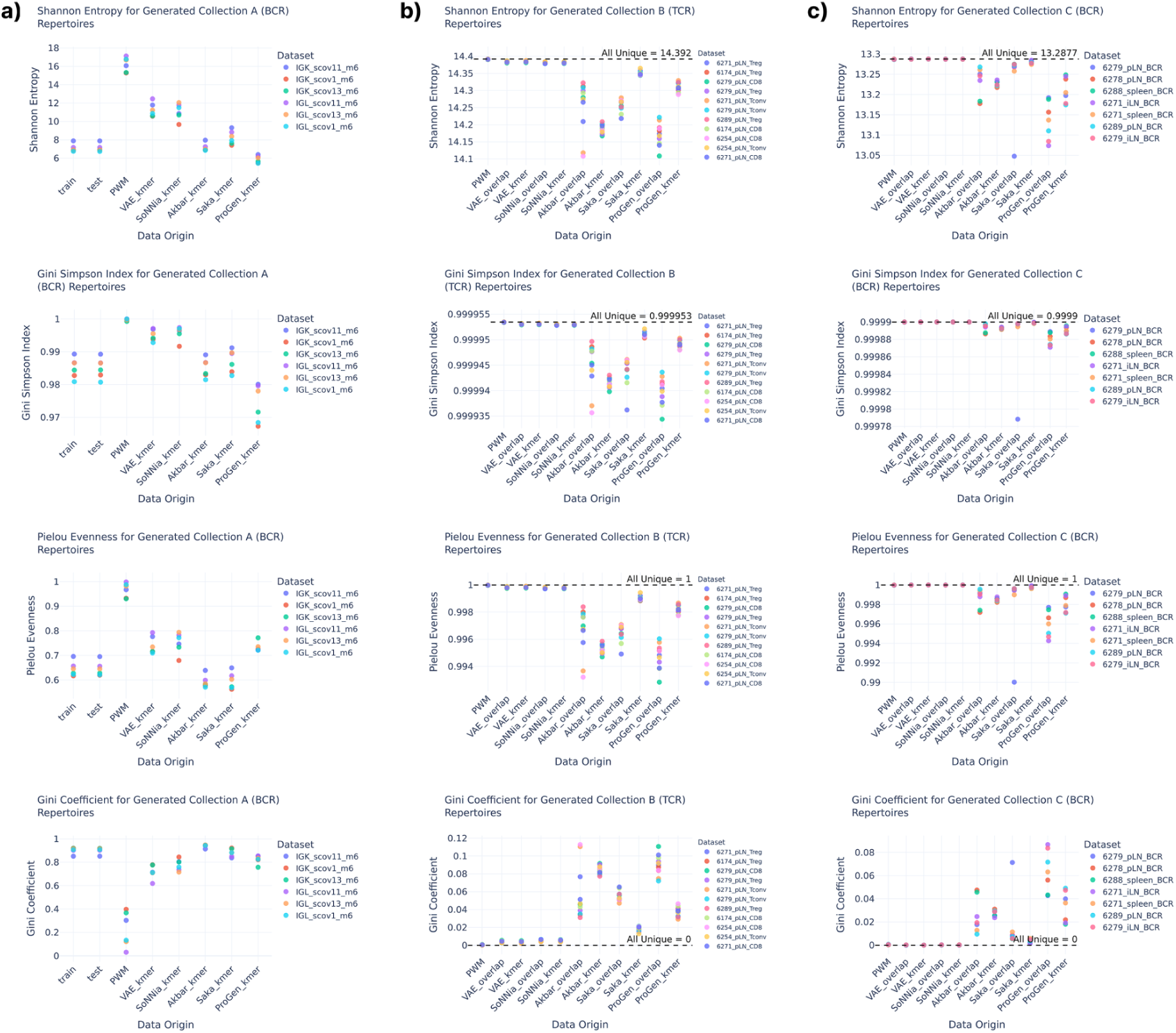
Clonal diversity scores per model and repertoire in the TCR and BCR collections with (a) clonally expanded cells and (b,c) unique sequences in reference repertoires. Diversity scores shown are Shannon entropy, Gini Simpson Index, Pielou Evenness, and Gini Coefficient. In (b,c), dashed reference lines indicate the score of the original datasets consisting of only unique sequence

**Figure S3:**
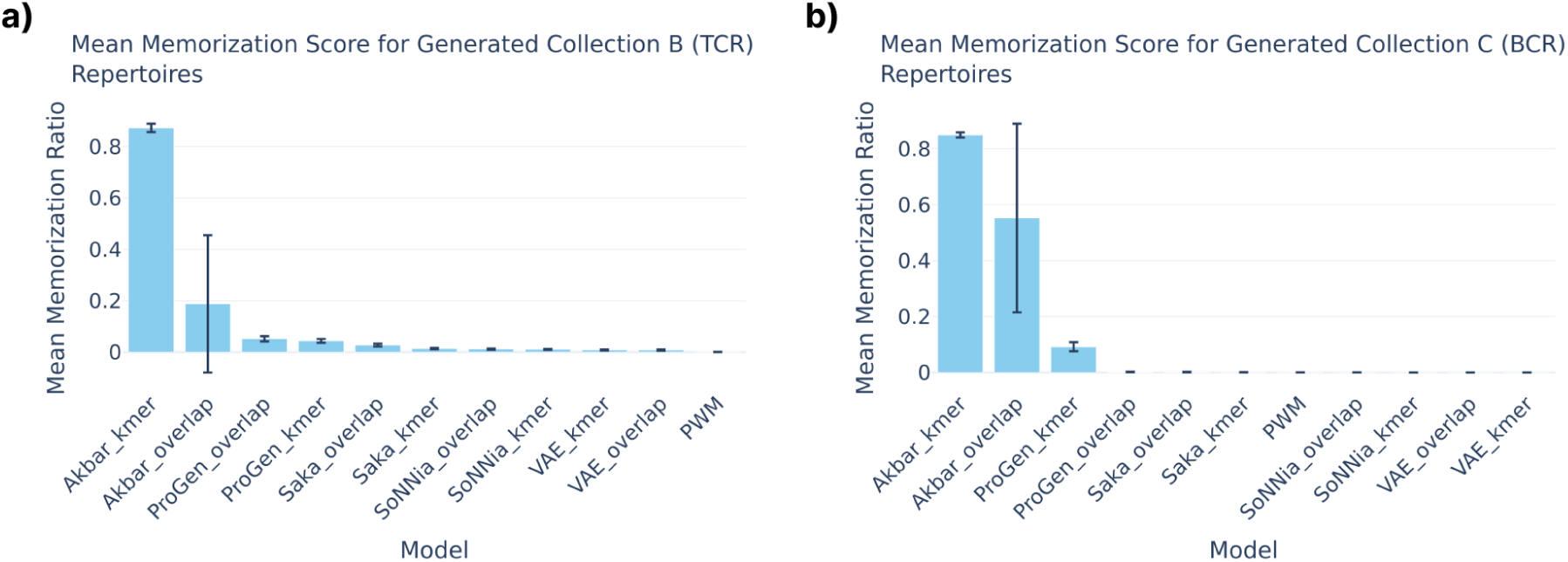
Memorization scores per generative model on TCR (a) and BCR (b) repertoires. Scores were averaged across batches of generated repertoires.

**Figure S4:**
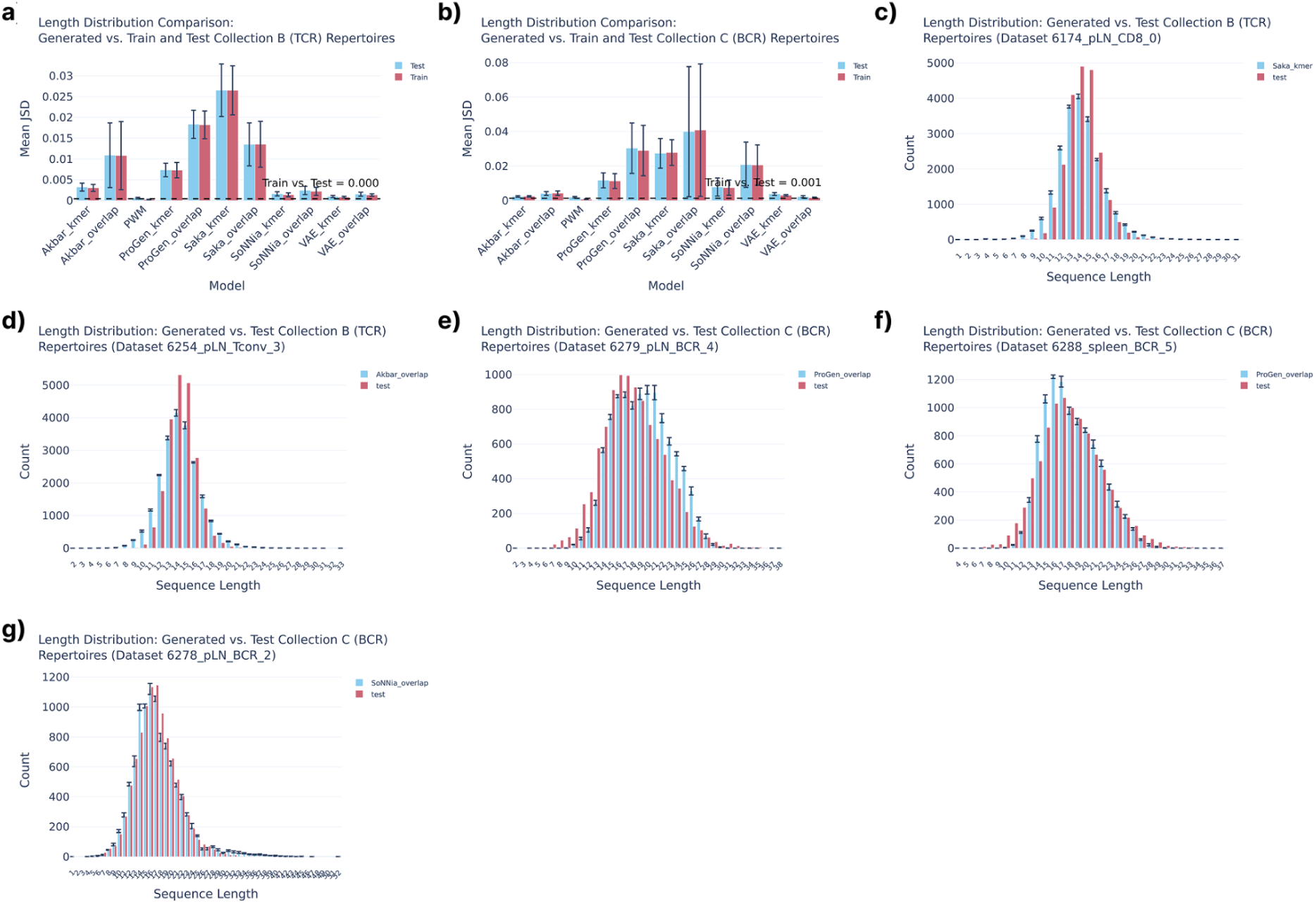
Comparisons of sequence length distributions for train, test, and generated TCR and BCR data (unique clones scenario). a,b) Jensen-Shannon divergence between train and generated, and test and generated sequence length distributions per model. Scores were averaged over subsets of generated repertoires. Dashed lines show corresponding reference scores computed between train and test sets. c-g) Sequence length distributions of exemplary models and subject-specific repertoires.

**Figure S5:**
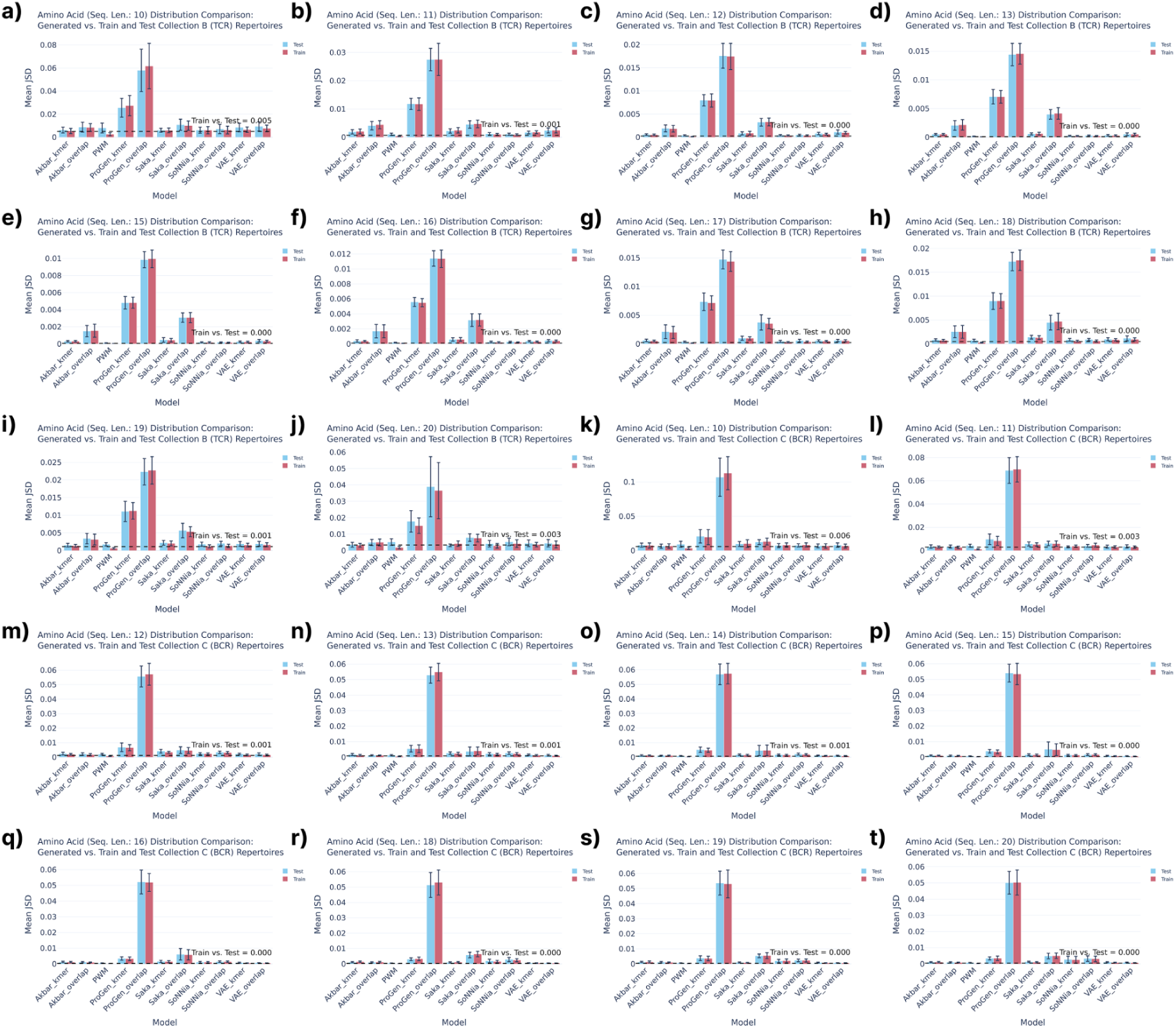
Comparisons of amino acid frequency distributions for train, test, and generated TCR (a-j) and BCR (k-t) repertoires (unique clones scenario). a-j) TCR distributions for sequence lengths 10-20 with exception of length 14. k-t) BCR distributions for sequence lengths 10-20 with exception of length 17. The recovery was measured by the Jensen-Shannon divergence with a lower score indicating higher distributional similarity. For each model, the average Jensen-Shannon divergence between train and generated, and test and generated was computed from the subsets of generated data. All dashed lines show corresponding reference scores computed between train and test sets.

**Figure S6:**
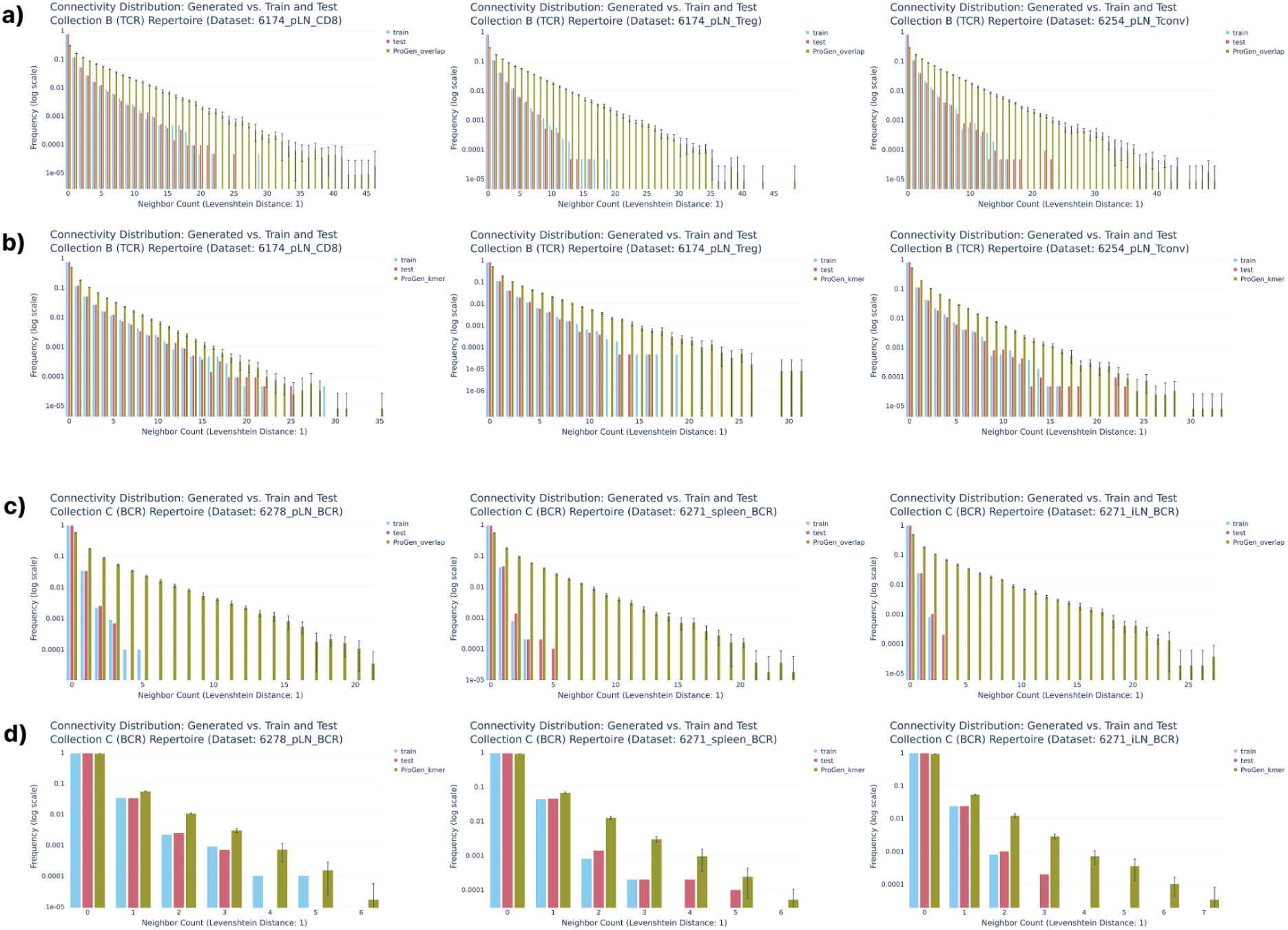
Comparison of sequence neighbor count distributions for train, test, and generated samples in (a,b) TCR and (c,d) BCR repertoires of three representative subjects from each data collection (A and B). ProGen models fine-tuned for innovation (a,c: ProGen_overlap) generally yielded higher neighbor counts than models fine-tuned to preserve low-dimensional characteristics (b,d: ProGen_kmer).

**Figure S7:**
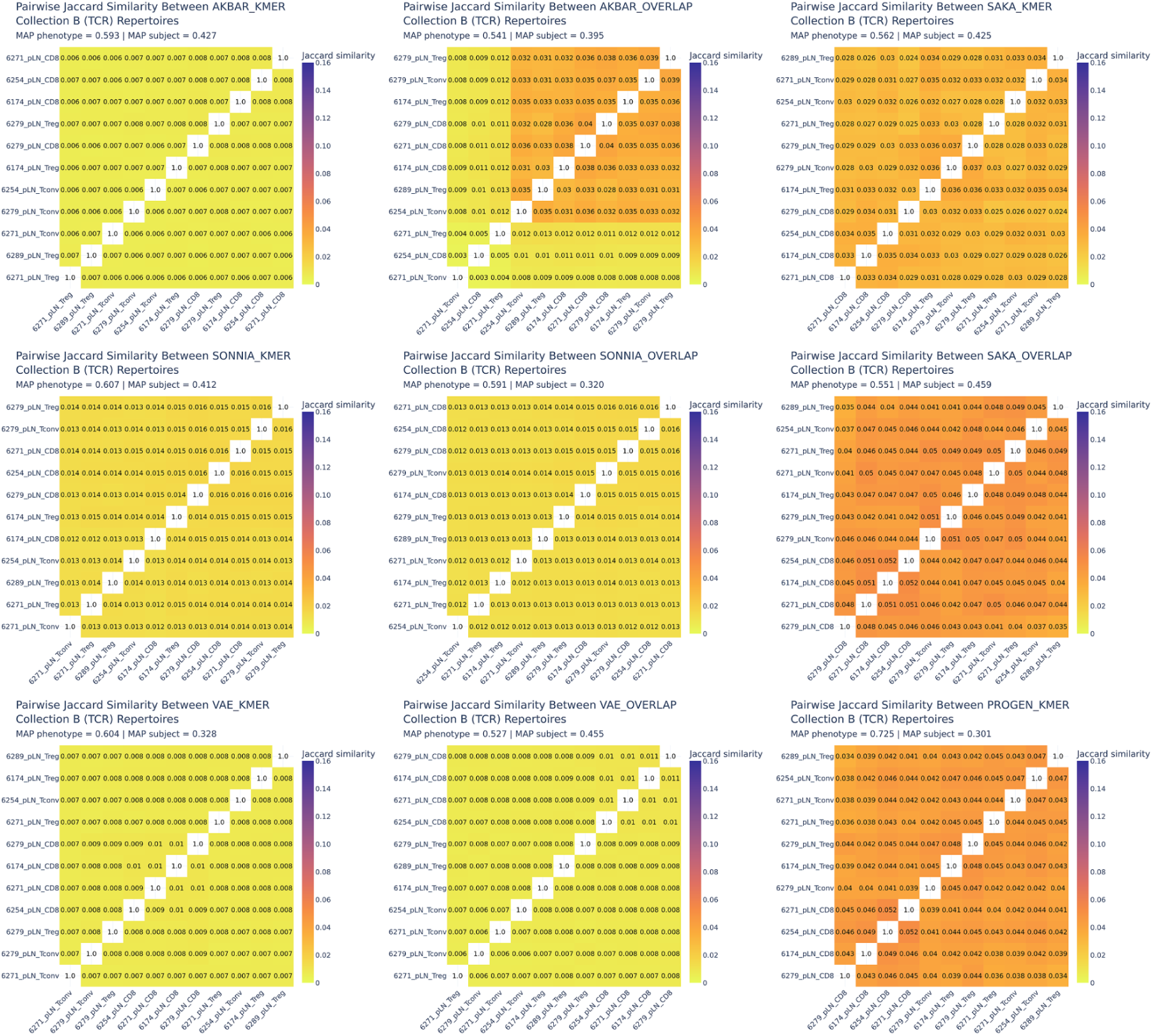
Hierarchical clustering of pairwise Jaccard similarity scores between generated TCR repertoires.

**Figure S8:**
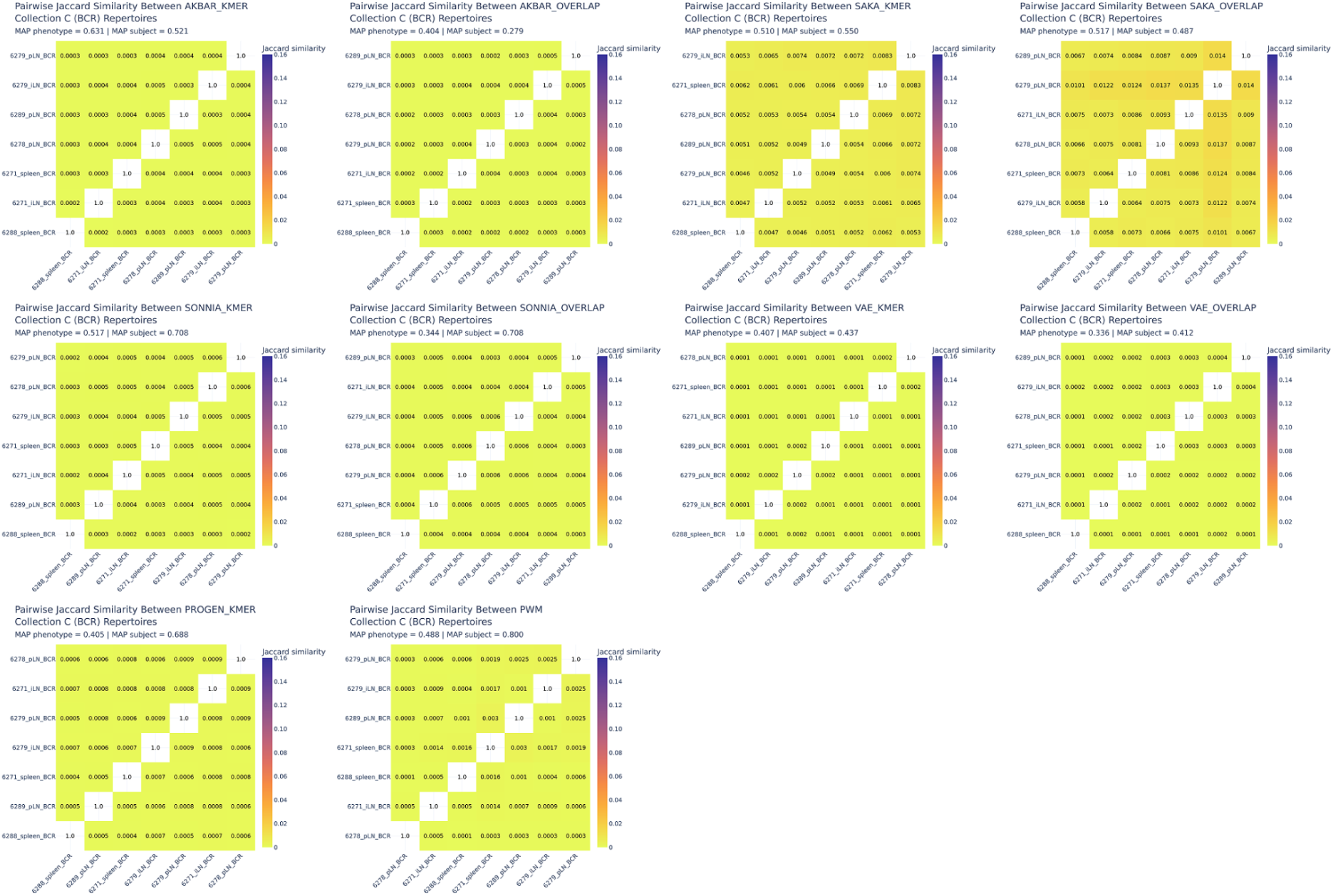
Hierarchical clustering of pairwise Jaccard similarity scores between generated BCR repertoires.

**Figure S9:**
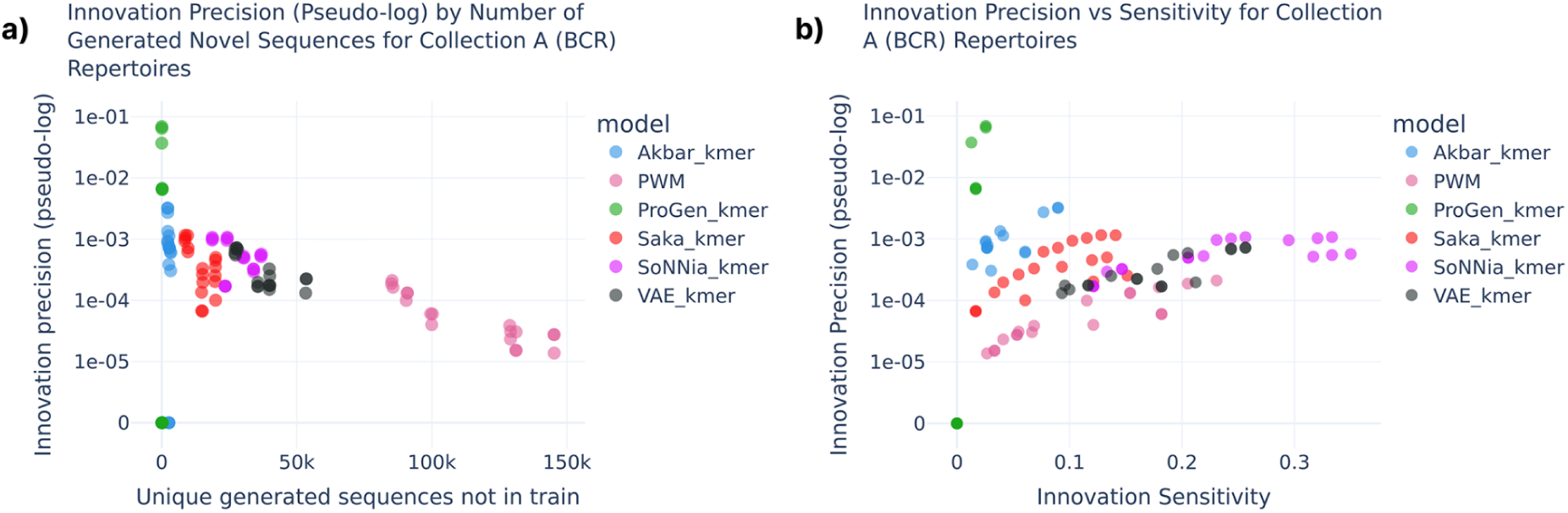
Innovation scores for models trained on clonally expanded repertoires (Collection A). a) Scatterplot of the number of unique generated sequences not present in the train repertoires vs. innovation precision for each generated subrepertoire, shown on symmetric log scale with threshold of 1e-05. b) Scatterplot of innovation sensitivity vs. innovation precision for each generated subrepertoire, shown on symmetric log scale with threshold of 1e-05.

## Supplementary Notes

### Model-specific factors limiting clone frequency recovery

As PWM models each position independently, it cannot capture co-occurrence patterns across positions. This causes it to miss precise sequence combinations of specific expanded clones, generating out-of-distribution sequences while failing to reproduce those dominating the training data (Figure S1c). The soNNia method combines a V(D)J background model with learned feature-specific selection factors, and the dissimilarity of the learned distribution could thereby originate from the more diverse background forcing the generated samples to contain fewer repetitions (Figure S1). VAE performed moderately better, yet it inherently aims to produce a continuous and diverse output distribution which naturally counteracts the more discrete nature of highly expanded exact sequences observed in the original repertoires (Figure S1e). Although the antibody language model (ProGen) memorized almost as much as Akbar, the model failed to reproduce sequence clone frequencies from the test samples (Figure S1f). Together with the lower diversity scores and higher neighbour counts (Figure S2, Figure S6), this indicates that ProGen produced certain regions of the distribution more densely, while other regions were underrepresented relative to the reference.

### Innovation in the clonal expansion scenario

Building on the precision-recall inspiration (27,28), we separated innovation quantification into two complementary scores to better handle the influence of sequence repetitions in clonally expanded repertoires. The innovation precision score measured the proportion of unique generated sequences (not found in the train sample) that had at least one direct sequence match in the test sample, while the innovation sensitivity score measured the proportion of unique unseen test sequences covered by at least one direct sequence match in the generated data. Restricting both metrics to unique generated sequences ensured that repeated sequences could not artificially inflate the innovation scores.

In the context of clonally expanded repertoires, where high-frequency sequences were present in both train and test, this experiment focused on recovery of the lower-frequency sequences unique to the test sample. As the models memorized train sequences to different degrees, the pools of unique generated sequences not seen during training varied in size.

Thus, innovation precision scores might have been inflated for models generating fewer unique sequences, as this reduces the denominator (Figure S9a). Although both ProGen and Akbar obtained the highest innovation precision scores, their performance was inconsistent across repertoires and their low numbers of unique generated sequences likely limited their ability to cover the unseen test sequence space, as reflected by the low innovation sensitivity scores (Figure S9b). When considering both innovation scores simultaneously, soNNia covered the largest proportion of test sequences while maintaining moderate precision. However, concluding on the models’ innovative capacities in the clonal expansion scenario is challenging as the numbers of test sequences not overlapping with the corresponding train samples were very few (fewer than 79 unique sequences depending on the repertoire).

### Recovery of sequence length distributions

For both TCR and BCR repertoires of unique sequences, the sequence length distribution was well captured by most models, with ProGen and Saka LSTM achieving poorer scores. (Figure S4a,b). Saka tended to overproduce shorter sequences (Figure S4c), while ProGen showed no consistent trend, sometimes overproducing short sequences and sometimes long ones (Figure S4e,f). Additionally, for TCR, we observed slightly poorer performance for Akbar LSTM optimized for the innovation criterion. The long error bars indicate that performance was inconsistent across repertoires (Figure S4a,b). In the case of BCR datasets, slightly poorer performance was obtained by SoNNia models, particularly when optimized for the innovation score. In these cases, the model sometimes generated distributions shifted toward shorter sequences, with a long tail toward longer sequences (Figure S4g).

## Supplementary Tables

**Table S1:** Overview of models and naming convention.

|  | Baseline | Variational Autoencoder | Long short-term memory |  | Selection model | Antibody language model |
| --- | --- | --- | --- | --- | --- | --- |
| Tuned method | PWM | VAE | Saka | Akbar | soNNia | ProGen2-OAS |
| Tuned to preserve k-mer distribution | NA | VAE_kmer | Saka_kmer | Akbar_kmer | soNNia_kmer | ProGen_kmer |
| Tuned for high innovation | NA | VAE_overlap | Saka_overlap | Akbar_overlap | soNNia_overlap | ProGen_overlap |

**Table S2:**
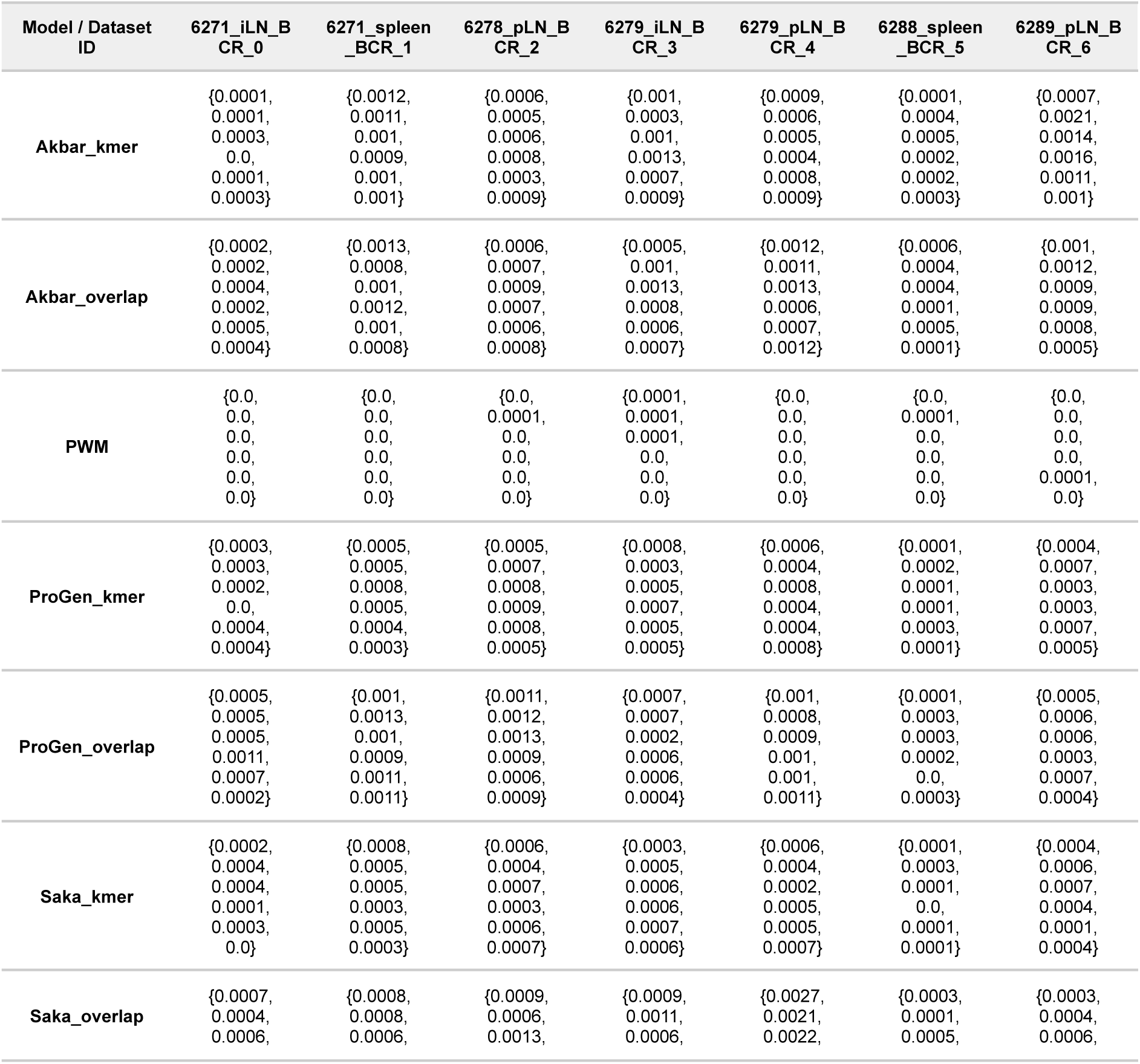
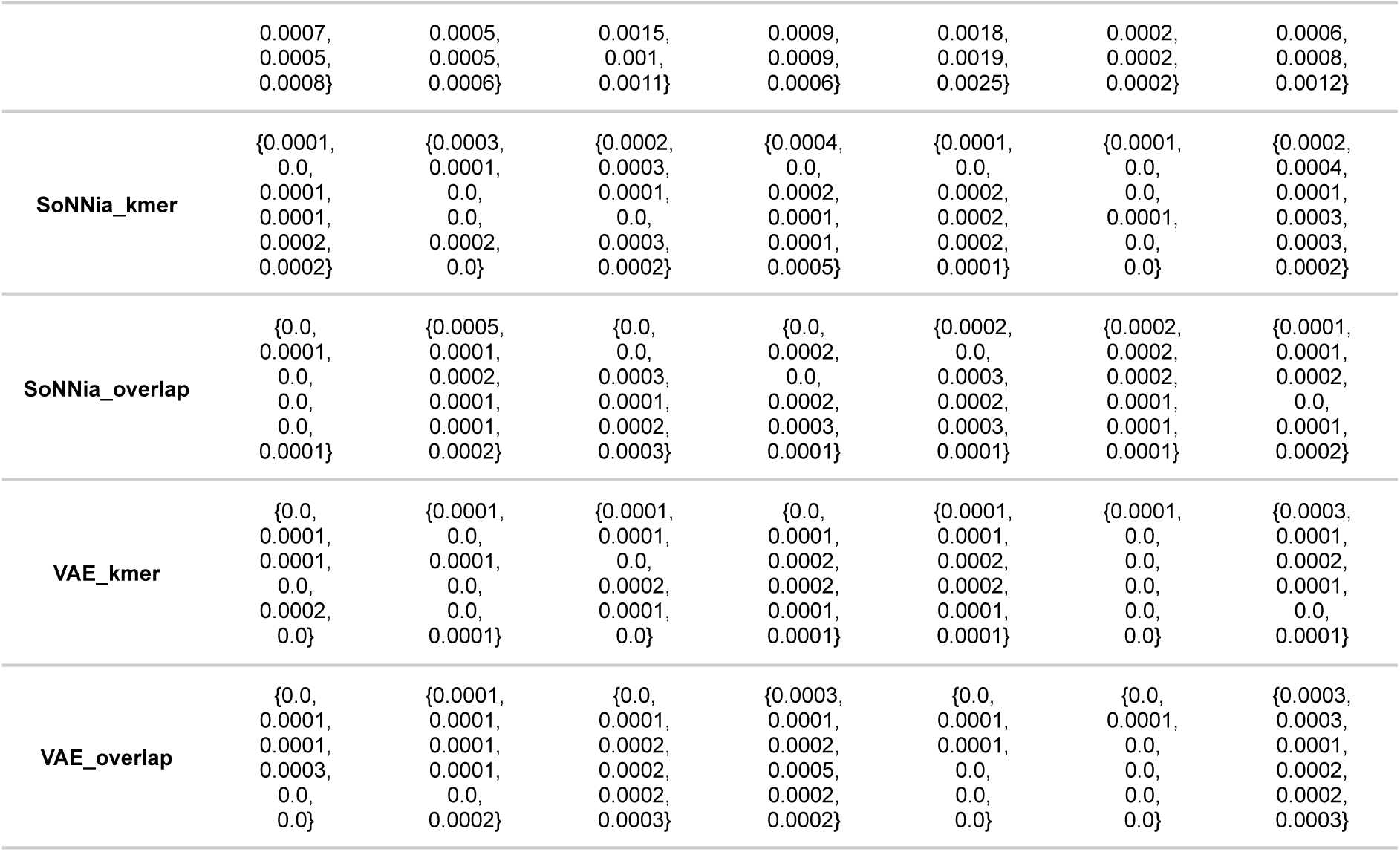
Innovation scores for collection C datasets.

**Table S3:** Hyperparameters of selected generative models.

| Hyperparameters | Clonal expansion scenario (BCR) |  |  |  |  | Unique clones scenario (TCR) |  |  |  |  |  |  |  |  |  | Unique clones scenario (BCR) |  |  |  |  |  |  |  |  |  |
| --- | --- | --- | --- | --- | --- | --- | --- | --- | --- | --- | --- | --- | --- | --- | --- | --- | --- | --- | --- | --- | --- | --- | --- | --- | --- |
|  | _kmer |  |  |  |  | _kmer / _overlap |  |  |  |  |  |  |  |  |  | _kmer / _overlap |  |  |  |  |  |  |  |  |  |
|  | Akbar | Saka | VAE | SoNNia | ProGen | Akbar |  | Saka |  | VAE |  | SoNNia |  | ProGen |  | Akbar |  | Saka |  | VAE |  | SoNNia |  | ProGen |  |
| Epochs | 20 | 100 | 1000 | 30 | 2 | 20 | 50 | 300 | 300 | 1000 | 1000 | 50 | 50 | 3 | 2 | 50 | 50 | 100 | 100 | 1000 | 500 | 30 | 100 | 3 | 1 |
| Batch size | 128 | 128 | 100 | 1000 | NA | 128 | 64 | 128 | 64 | 100 | 100 | 500 | 1000 | NA | NA | 128 | 128 | 128 | 64 | 100 | 100 | 1000 | 1000 | NA | NA |
| Hidden size | 512 | 128 | NA | NA | NA | 1024 | 1024 | 64 | 128 | NA | NA | NA | NA | NA | NA | 512 | 1024 | 128 | 64 | NA | NA | NA | NA | NA | NA |
| Learning rate | 0.001 | 0.01 | 0.001 | NA | 4E-05 | 0.001 | 0.001 | 0.01 | 0.01 | 0.001 | 0.001 | NA | NA | 4E-05 | 4E-05 | 0.001 | 0.001 | 0.01 | 0.01 | 0.001 | 0.001 | NA | NA | 4E-05 | 4E-05 |
| Embedding size | 256 | 256 | NA | NA | NA | 256 | 256 | 256 | 256 | NA | NA | NA | NA | NA | NA | 256 | 256 | 256 | 256 | NA | NA | NA | NA | NA | NA |
| Temperature | 1 | 1 | NA | NA | 1 | 1 | 0.8 | 1 | 0.8 | NA | NA | NA | NA | 1 | 0.8 | 1 | 0.8 | 1 | 0.8 | NA | NA | NA | NA | 1 | 0.8 |
| Number of layers | 3 | 2 | NA | NA | NA | 2 | 3 | 3 | 3 | NA | NA | NA | NA | NA | NA | 2 | 3 | 2 | 3 | NA | NA | NA | NA | NA | NA |
| Number of frozen layers | NA | NA | NA | NA | 22 | NA | NA | NA | NA | NA | NA | NA | NA | 22 | 22 | NA | NA | NA | NA | NA | NA | NA | NA | 19 | 22 |
| Top p | NA | NA | NA | NA | 0.9 | NA | NA | NA | NA | NA | NA | NA | NA | 0.9 | 0.9 | NA | NA | NA | NA | NA | NA | NA | NA | 0.9 | 0.7 |
| beta | NA | NA | 0.75 | NA | NA | NA | NA | NA | NA | 0.75 | 0.75 | NA | NA | NA | NA | NA | NA | NA | NA | 0.75 | 0.75 | NA | NA | NA | NA |
| Warm up epochs | NA | NA | 20 | NA | NA | NA | NA | NA | NA | 20 | 20 | NA | NA | NA | NA | NA | NA | NA | NA | 20 | 20 | NA | NA | NA | NA |
| Pretrains | NA | NA | 10 | NA | NA | NA | NA | NA | NA | 10 | 10 | NA | NA | NA | NA | NA | NA | NA | NA | 10 | 10 | NA | NA | NA | NA |
| Patience | NA | NA | 20 | NA | NA | NA | NA | NA | NA | 20 | 20 | NA | NA | NA | NA | NA | NA | NA | NA | 20 | 20 | NA | NA | NA | NA |
| Latent dimension | NA | NA | 30 | NA | NA | NA | NA | NA | NA | 30 | 20 | NA | NA | NA | NA | NA | NA | NA | NA | 20 | 30 | NA | NA | NA | NA |
| Linear node count | NA | NA | 125 | NA | NA | NA | NA | NA | NA | 125 | 25 | NA | NA | NA | NA | NA | NA | NA | NA | 125 | 25 | NA | NA | NA | NA |
| N gen seqs | NA | NA | NA | 1M | NA | NA | NA | NA | NA | NA | NA | 200K | 1M | NA | NA | NA | NA | NA | NA | NA | NA | 500K | 1M | NA | NA |

**Table S4:**
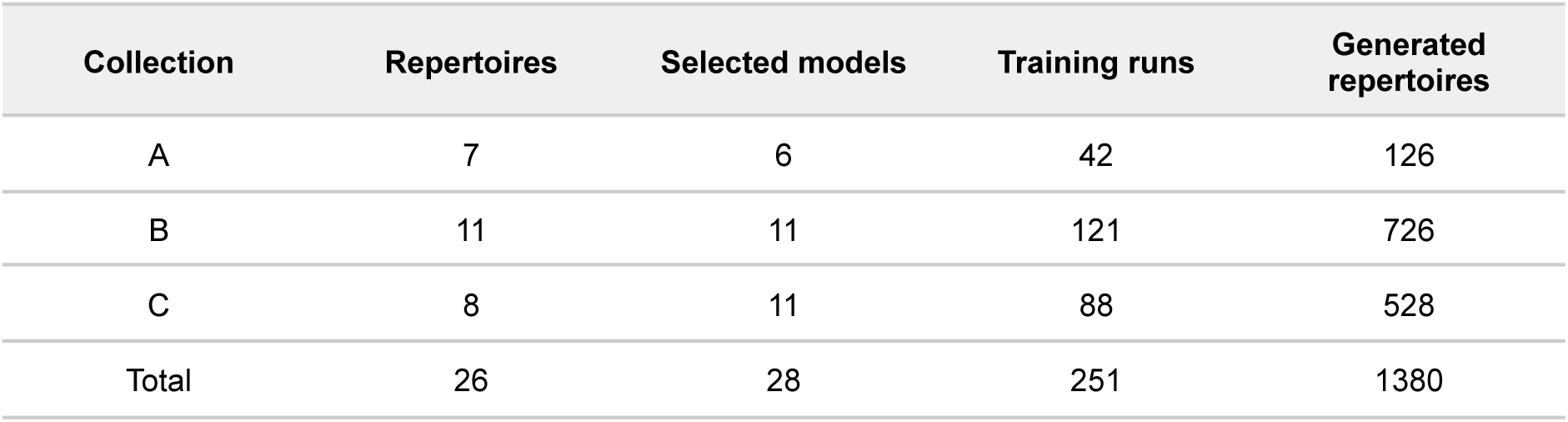
Computational overview of model training and evaluation. For each data collection, the table reports the number of repertoires, selected models, model training runs, and generated repertoires. Selected models correspond to the final hyperparameter configurations selected after model tuning.

| Collection | Repertoires | Selected models | Training runs | Generated repertoires |
| --- | --- | --- | --- | --- |
| A | 7 | 6 | 42 | 126 |
| B | 11 | 11 | 121 | 726 |
| C | 8 | 11 | 88 | 528 |
| Total | 26 | 28 | 251 | 1380 |

**Table S5:** Hyperparameter tuning was carried out independently on one held-out representative repertoire from each collection. The table reports the number of tested hyperparameter configurations, the corresponding number of trained models across all collections, and the number of generated repertoires used for model selection. Each tested configuration was trained independently for each collection.

| Model | Configurations tested | Total models trained for tuning (across all collections) | Generated repertoires for tuning (A) | Generated repertoires for tuning (B) | Generated repertoires for tuning (C) | Total generated repertoires for tuning |
| --- | --- | --- | --- | --- | --- | --- |
| Akbar | 32 | 96 | 96 | 192 | 192 | 480 |
| Saka | 32 | 96 | 96 | 192 | 192 | 480 |
| VAE | 24 | 72 | 72 | 144 | 144 | 360 |
| SoNNia | 24 | 72 | 72 | 144 | 144 | 360 |
| ProGen | 36 | 108 | 108 | 216 | 216 | 540 |
| Total | 148 | 444 | 444 | 888 | 888 | 2220 |

**Table S6:** Overview of repertoire dataset collections A, B, and C. (A) clonally expanded BCR repertoires with sequence frequencies accounted for through unique molecular identifiers (UMIs). Repertoires originate from blood samples of SARS-CoV-2 patients 6 months after disease onset (52). (B) TCR repertoires with only unique clones retained. (C) BCR repertoires with only unique clones retained.

| Collection | Phenotype | Tissue | Locus | Cell-subset | Subject id | Sample id | Used for hyper-parameter tuning | Sequences sampled for train & test | Unique sequences in sampled train | Unique sequences in sampled test |
| --- | --- | --- | --- | --- | --- | --- | --- | --- | --- | --- |
| A | IGL | peripheral blood | IGL | mature B cell | S-Cov1 | S-CoV1_M6_Sort1 | No | 150.000 | 1842 | 1841 |
| A | IGL | peripheral blood | IGL | mature B cell | S-Cov11 | S-CoV11_M6_Sort1 | No | 150.000 | 1936 | 1955 |
| A | IGL | peripheral blood | IGL | mature B cell | S-Cov13 | S-CoV13_M6_Sort1 | No | 150.000 | 1670 | 1673 |
| A | IGK | peripheral blood | IGK | mature B cell | S-Cov1 | S-CoV1_M6_Sort1 | No | 150.000 | 2251 | 2220 |
| A | IGK | peripheral blood | IGK | mature B cell | S-Cov11 | S-CoV11_M6_Sort1 | No | 150.000 | 2595 | 2600 |
| A | IGK | peripheral blood | IGK | mature B cell | S-Cov13 | S-CoV13_M6_Sort1 | No | 150.000 | 2092 | 2091 |
| A | IGK | peripheral | IGK | mature B | S-Cov10 | S-CoV10_M6_S | Yes | 150.000 | 1789 | 1777 |

|  |  | blood |  | cell |  | ort1 |  |  |  |  |
| --- | --- | --- | --- | --- | --- | --- | --- | --- | --- | --- |
| B | CD8+ | pancreatic lymph node | TRB | CD8+ | 6174 | 6174_pancreatic lymph node_CD8+ T cell | No | 21.500 | 21.500 | 21.500 |
| B | CD8+ | pancreatic lymph node | TRB | CD8+ | 6271 | 6271_pancreatic lymph node_CD8+ T cell | No | 21.500 | 21.500 | 21.500 |
| B | CD8+ | pancreatic lymph node | TRB | CD8+ | 6279 | 6279_pancreatic lymph node_CD8+ T cell | No | 21.500 | 21.500 | 21.500 |
| B | CD8+ | pancreatic lymph node | TRB | CD8+ | 6254 | 6254_pancreatic lymph node_CD8+ T cell | No | 21.500 | 21.500 | 21.500 |
| B | CD4+ Tconv | pancreatic lymph node | TRB | CD4+ Tconv | 6271 | 6271_pancreatic lymph node_CD4+CD127+ Tconv | No | 21.500 | 21.500 | 21.500 |
| B | CD4+ Tconv | pancreatic lymph node | TRB | CD4+ Tconv | 6254 | 6254_pancreatic lymph node_CD4+CD127+ Tconv | No | 21.500 | 21.500 | 21.500 |
| B | CD4+ Tconv | pancreatic lymph node | TRB | CD4+ Tconv | 6279 | 6279_pancreatic lymph node_CD4+CD127+ Tconv | No | 21.500 | 21.500 | 21.500 |
| B | CD4+ Treg | pancreatic lymph node | TRB | CD4+ Treg | 6174 | 6174_pancreatic lymph node_CD4+CD127-CD25+ Treg | No | 21.500 | 21.500 | 21.500 |
| B | CD4+ Treg | pancreatic lymph node | TRB | CD4+ Treg | 6289 | 6289_pancreatic lymph node_CD4+CD127-CD25+ Treg | No | 21.500 | 21.500 | 21.500 |
| B | CD4+ Treg | pancreatic lymph node | TRB | CD4+ Treg | 6271 | 6271_pancreatic lymph node_CD4+CD127-CD25+ Treg | No | 21.500 | 21.500 | 21.500 |
| B | CD4+ Treg | pancreatic lymph node | TRB | CD4+ Treg | 6279 | 6279_pancreatic lymph node_CD4+CD127-CD25+ Treg | No | 21.500 | 21.500 | 21.500 |
| B | CD4+ Treg | pancreatic lymph node | TRB | CD4+ Treg | 6288 | 6288_pancreatic lymph node_CD4+CD127-CD25+ Treg | Yes | 21.500 | 21.500 | 21.500 |
| C | BCR spleen | spleen | IGH | B cell | 6288 | 6288_spleen_CD19+_B_cell | No | 10.000 | 10.000 | 10.000 |
| C | BCR spleen | spleen | IGH | B cell | 6271 | 6271_spleen_CD19+_B_cell | No | 10.000 | 10.000 | 10.000 |
| C | BCR spleen | spleen | IGH | B cell | 6279 | 6279_spleen_CD19+_B_cell | No | 10.000 | 10.000 | 10.000 |
| C | BCR iLN | irrelevant lymph node | IGH | B cell | 6271 | 6271_irrelevant_lymph_node_CD19+_B_cell | No | 10.000 | 10.000 | 10.000 |
| C | BCR iLN | irrelevant lymph node | IGH | B cell | 6279 | 6279_irrelevant_lymph_node_CD19+_B_cell | No | 10.000 | 10.000 | 10.000 |
| C | BCR pLN | pancreatic lymph node | IGH | B-cell | 6278 | 6278_pancreatic_lymph_node_CD19+ B cell | No | 10.000 | 10.000 | 10.000 |
| C | BCR pLN | pancreatic lymph node | IGH | B-cell | 6279 | 6279_pancreatic_lymph_node_CD19+ B cell | No | 10.000 | 10.000 | 10.000 |
| C | BCR pLN | pancreatic lymph node | IGH | B cell | 6289 | 6289_pancreatic_lymph_node_CD19+_B_cell | No | 10.000 | 10.000 | 10.000 |
| C | BCR pLN | pancreatic lymph node | IGH | B cell | 6271 | 6271_pancreatic_lymph_node_CD19+ B cell | Yes | 10.000 | 10.000 | 10.000 |

**Table S7:** Summary table of repertoire dataset collections A, B, and C.

| Collection | Tissue | Cell type | Number of repertoires | Weighted by UMI counts | Number of sequences sampled for train & test | Avg. num. unique train sequences after sampling | Avg. num. unique test sequences after sampling | Number of sampled batches |
| --- | --- | --- | --- | --- | --- | --- | --- | --- |
| A | Peripheral blood | IGL | 3 | Yes | 150.000 | 1816 | 1823 | 3 |
| A | Peripheral blood | IGK | 4 | Yes | 150.000 | 2182 | 2172 | 3 |
| B | Pancreatic lymph node | TRB CD8+ | 4 | No | 21.500 | 21.500 | 21.500 | 6 |
| B | Pancreatic lymph node | TRB CD4+ Tconv | 3 | No | 21.500 | 21.500 | 21.500 | 6 |
| B | Pancreatic lymph node | TRB CD4+ Treg | 5 | No | 21.500 | 21.500 | 21.500 | 6 |
| C | Spleen | IGH | 3 | No | 10.000 | 10.000 | 10.000 | 6 |
| C | Pancreatic lymph node | IGH | 2 | No | 10.000 | 10.000 | 10.000 | 6 |
| C | Irrelevant lymph node | IGH | 3 | No | 10.000 | 10.000 | 10.000 | 6 |

